# Fetal bovine serum-triggered Titan cell formation and growth inhibition are unique to the *Cryptococcus* species complex

**DOI:** 10.1101/435842

**Authors:** Mariusz Dylag, Rodney Colon-Reyes, Lukasz Kozubowski

## Abstract

In the genus *Cryptococcus*, *C. neoformans* and *C. gattii* stand out by the number of virulence factors that allowed those fungi to achieve evolutionary success as pathogens. Among the factors contributing to cryptococcosis is a peculiar morphological transition to form giant (Titan) cells. Formation of Titans has been described *in vitro*. However, it remains unclear whether non-*C. neoformans*/non-*C. gattii* species are capable of titanisation. Based on a survey of several basidiomycetous yeasts, we propose that titanisation is unique to *C. neoformans* and *C. gattii*. In addition, we find that under *in vitro* conditions that induce titanisation, fetal bovine serum (FBS) possesses activity that inhibits growth of *C. gattii* and kills *C. neoformans* if incubation is conducted in the absence of 5% CO_2_. Moreover, lowering of pH or addition of an antioxidant, rescues growth in the presence of FBS and allows titanisation even in the absence of 5% CO_2_. Strikingly, acapsular mutants, *cap10*Δ and *cap59*Δ, show complete or partial reduction of titanisation, respectively and a partial resistance to growth inhibition imposed by the FBS. Our data implicate titanisation as a unique feature of *C. neoformans* and *C. gattii* and provide novel insights about the nature of this unusual morphological transition.

## INTRODUCTION

Basidiomycetous yeasts are an ecologically heterogeneous group of fungi that includes both saprophytic as well as pathogenic species. A significant fraction of the latter group comprises of human and animal pathogens, including genera of *Cryptococcus*, *Malassezia*, *Rhodotorula* and *Trichosporon* ^1^. In particular, two species belonging to the *Cryptococcus* genus (formerly Filobasidiella), *C. neoformans* and *C. gattii* are the etiological agents of fatal systemic mycoses. In recent years, nearly one million cases of cryptococcal meningitis occur annually resulting in over 600,000 deaths globally ^2^. Moreover, *cryptococcosis* is responsible for 15-17% AIDS-related deaths on a global scale ^3,4^. While *C. neoformans* can cause mainly opportunistic infections in immunocompromised patients, *C. gattii* is capable of infecting also immunocompetent individuals ^5–8^. Moreover, it is well established that among *C. neoformans*, serotype A (varietas *grubii*) is responsible for the majority of cryptococcal infections, while serotype D (varietas *neoformans*) is less common ^7^. Among *Cryptococcus* species other than *C. neoformans* and *C. gattii*, the following species have been described as causing occasional infections in humans: *C. laurentii* ^9–11^, *C. albidus* ^9,12,13^, *C. curvatus* ^14,15^, *C. uniguttulatus* ^16^, and *C. adeliensis* ^17^. Such casuistic infections, reviewed in literature most recently by Smith et al. ^18^ can be also systemic in case of strains able to grow at 37°C. What makes those selected *Cryptococcus* species capable of infecting humans is an important question, the answer to which remains incomplete. Among the best-described cryptococcal characteristics necessary for pathogenicity are the ability to proliferate at 37°C, melanisation, formation of capsule, and the capability of hydrolyzing urea ^8,19^. Most of the above-mentioned “virulence factors” can be observed together only in case of *C. neoformans* and *C. gattii*, what could potentially explain the basis of their evolutionary success as pathogens.

Dimorphism is one of the common features of human fungal pathogens ^20,21^. A well-documented example of a dimorphic switching critical for pathogenicity is yeast to hypha transition characteristic for dimorphic fungi (clustered within five genera) and *Candida albicans* ^20, 21^. Morphological transition of *C. neoformans* to form enlarged cells termed Titan cells is a particularly striking manifestation of a perfect adaptation to evade the mammalian immune system and enhance dissemination in the host ^22^. Titan cells are resistant to phagocytosis due to over ten-time enlargement of the cell body ^22–25^. Moreover, it has been demonstrated that Titan cells and aneuploid daughter cells of Titans show increased resistance to many physico-chemical factors and some drugs commonly used in the therapy of cryptococcosis ^26^. However, titanisation may not be required for pathogenicity as some clinical isolates of *C. neoformans* are presumably not capable of undergoing this morphological transition ^27–29^. Therefore, it remains unclear to what extend titanisation is important for cryptococcosis and whether this striking characteristic is unique to pathogenic *Cryptococcus* species, especially since titanisation has never been studied in species other than *C. neoformans* and *C. gattii*.

Recent investigations led to considerable advances in our understanding of the nature of titanisation, by uncovering external stimuli that are sufficient and genes that are essential for this morphological transition ^27–29^. Two well-documented pathways involved in titanisation are the cAMP-mediated signaling (dependent on Gpr4/Gpr5 receptors, adenylyl cyclase Cac1, Pka1 kinase, and the transcription factor Rim101) and the mating pathway ^30–32^. Until recently the progress in identifying more regulators has been hampered by the lack of a suitable *in vitro* system. However, recently published work from several laboratories delivered new opportunities towards thorough understanding of titanisation by developing *in vitro* conditions capable of stimulating this morphological change ^27–29^. This breakthrough research led to identification of novel positive (Gat201, Ada2, Cap59, Cap60, Ric8, Sgf29, Lmp1) and negative (Usv101, Pkr1, Tsp2) regulators that were important for titanisation under specific conditions ^27,29^. These studies also led to an overarching conclusion that titanisation can be stimulated by a variety of external signals, including CO_2_, hypoxia, exposure to serum (specifically two serum components, phospholipids and bacterial peptidoglycan), and quorum sensing ^27–29^. Importantly, these novel protocols to induce titanisation *in vitro* have provided an opportunity to test to what extent is the ability to form Titans conserved in other *Cryptococcus* species.

Based on a survey of 23 strains that represent 9 non-*neoformans* species, we postulate that *C. neoformans* and *C. gattii* are unique among other basidiomycetous yeasts with regard to the ability to form *bona fide* Titan cells in the presence of fetal bovine serum. Our work provides novel insights with respect to external factors that are necessary, sufficient, or inhibitory towards titanisation. We uncover that fetal bovine serum contains both inhibitory and stimulatory factors and a balance between these factors determines the ability to form Titan cells. The inhibitory factor affects growth through increase of ROS. In contrast the stimulatory factor triggers titanisation likely through cAMP pathway, consistent with previous findings. Furthermore, exposure to 5% CO_2_ counteracts the inhibitory effect of serum and further stimulates Titan cell formation. Strikingly, *C. gattii* and even more significantly *C. neoformans* var. *grubii* are particularly sensitive towards the inhibitory effect of serum as compared to other non-*neoformans Cryptococcus* species. We also demonstrate that deletion of genes involved in capsule formation, *CAP10* and *CAP59*, have differential effect on titanisation; *CAP10* is essential and *CAP59* dispensable for this morphogenetic transition. Our data suggest that *C. neoformans* and *C. gattii* have evolved as unique species capable of undergoing titanisation under human host conditions.

## RESULTS

### *C. neoformans* and *C. gattii* are unique in their ability to undergo titanisation in the presence of fetal bovine serum (FBS)

Three *in vitro* protocols to induce formation of *bona fide* Titan cells in *C. neoformans* have been published recently ^27–29^. In all three cases putative Titan cells formed that met established criteria characteristic of cell gigantism observed originally *in vivo* during murine infection caused by *C. neoformans* var. *grubii* ^25,31^. Typically *C. neoformans* var. *grubii* cells range in size from 4 to 6 μm, whereas the size larger than 10 μm has been considered as indicative of titanisation. Other key features of Titan cells are thicker cell wall, single large vacuole and increased ploidy within a single nucleus ^25,31,33^. Furthermore, Titans can produce daughters of “normal” size, which can be haploid or frequently aneuploid ^26^.

We utilized a method based on the protocol described by Dambuza et al. to test nine non-*neoformans Cryptococcus* species for their ability to undergo titanisation *in vitro* ^28^ The following species were tested: *C. albidus* (2 strains), *C. aspenensis* (4 strains), *C. curvatus* (2 strains), *C. gattii* (2 strains), *C. kuetzingii* (1 strain), *C. laurentii* (5 strains), *C. terrestris* (4 strains), *C. terreus* (1 strain), and *C. uniguttulatus* (2 strains). To include other basidiomycetous yeasts outside of the *Cryptococcus* genus, we tested two species of *Malassezia*: *M. furfur* (1 strain) and *M. sympodialis* (1 strain). We also included one reference strain of *Saccharomyces cerevisiae* as a representative of ascomycetous yeasts. As an additional control, we included ten *C. neoformans* var. *grubii* environmental strains and two strains of *C. neoformans* var. *neoformans* (all strains tested are listed in Table S1).

As titanisation protocol introduced by Dambuza et al. recommended incubation at 37°C, we first analyzed how temperature affected growth of the strains included in our study. We reasoned that the ideal temperature for titanisation is close to but not above the maximum temperature a given strain tolerates. While *C. neoformans* was capable of proliferating at 37°C, as reported previously, the majority of non-*neoformans* species could not grow at this temperature (Table S1). Based on the fact that the majority of strains could not proliferate at 37°C and grew at 30°C, we decided to utilize 30°C when testing titanisation and compare to 37°C for all the strains tested.

We estimated the percentage of Titan cells in the population based on a visual inspection of cells after 48 h of incubation under titanisation conditions. Specifically, we defined Titan cells as cells with diameter of close to or larger than 10 μm, with characteristic single large vacuole, and a thick cell wall as judged based on differential interference contrast (DIC) microscopy. As we decided to test Titan cell formation at 30°C, it was important to assess whether lowering the temperature to 30°C affects titanisation of *C. neoformans* var. *grubii* and *C. gattii*. Consistent with previous findings, *C. neoformans* var. *grubii* strain H99 formed Titan cells after 48 hours of incubation in 10% fetal bovine serum (FBS) under 5% CO_2_ at 37°C (Figure 1, Table S2). We estimated that cultures of eleven out of twelve strains of *C. neoformans* var *grubii*, tested at 37°C (one of the environmental isolates that has been collected in North Carolina, strain BS001, did not titanize), contained an average of 9.7 +/− 1.5% Titan cells after 48 hours (Table S2). Importantly all these strains proliferated as the number of cells within 48 hours increased between nearly 500 to over 1000 times from the initial inoculum of ~10^3^ cells (Table S2). Dambuza et al. have reported that under conditions similar to ours, *C. gattii* strain R265 failed to form Titans, whereas conditions established by Trevijano-Contador et al. led to titanisation of this strain ^27,28^. We considered that some of the other strains of *C. gattii* form Titans under our conditions. Consistent with this possibility, an environmental isolate of *C. gattii* (WM276) as well as the clinical strain R265 formed Titan cells under our conditions (Figure 1, Table S2). Notably, the percentage of Titan cells and the average size of cells were higher for *C. gattii* as compared to *C. neoformans* var. *grubii* (Figure 1, Table S2). However, in contrast to *C. neoformans* var. *grubii*, *C. gattii* strains did not proliferate significantly within 48 hours as the number of cells only increased 3-5 times (Table S2). *C. neoformans* var. *neoformans* (serotype D) strain JEC21 proliferated robustly but did not form Titan cells (Table S2). Strain JEC20 of the same serotype was strongly inhibited for growth and also did not exhibit titanisation under these conditions (Table S2).

**Figure 1.**
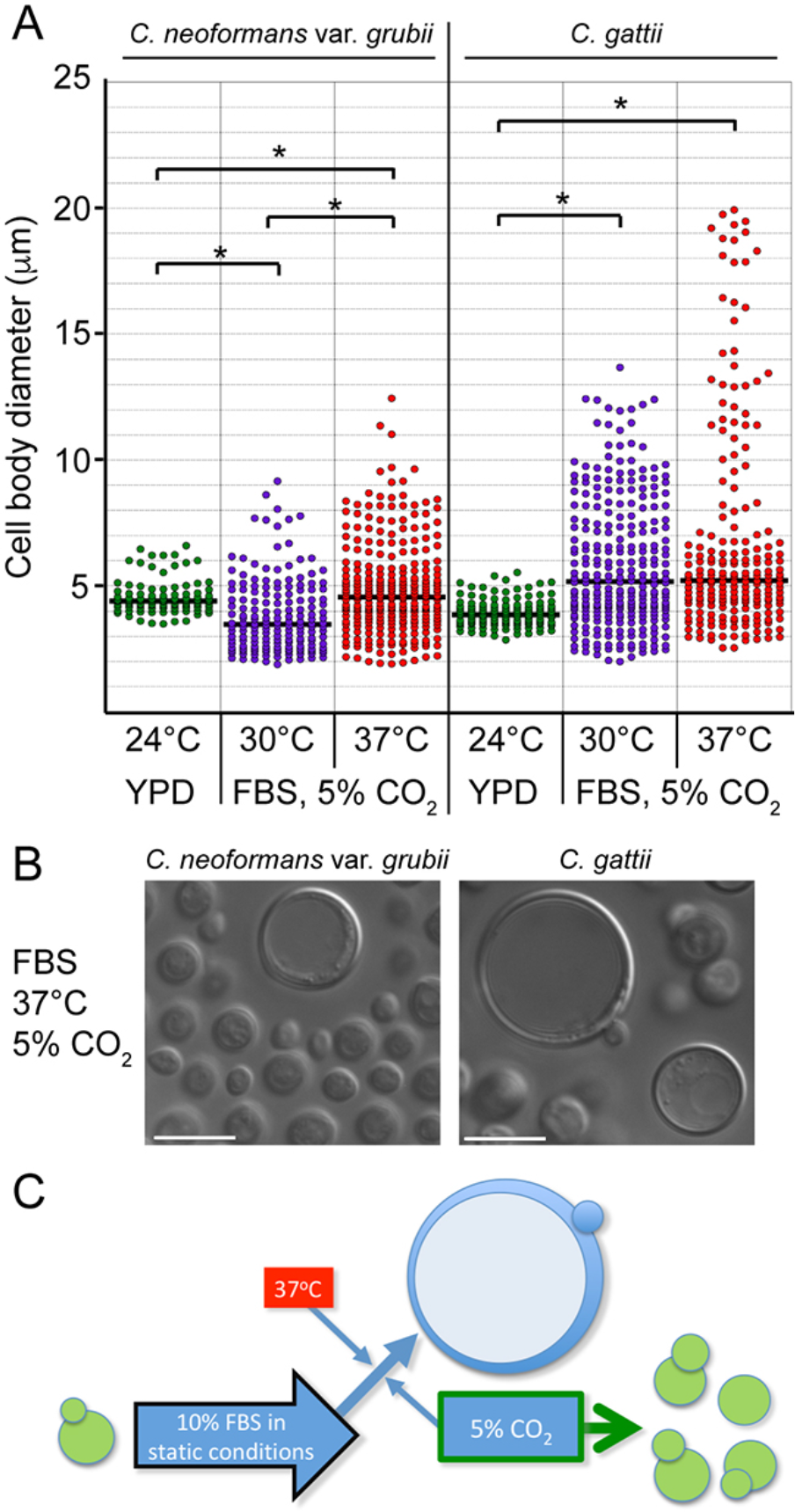
The effect of temperature on the induction of Titan cells in *C. neoformans* var.*grubii* strain H99 and *C. gattii* strain R265. Cells were subject to titanisation protocol ^28^ at either 30 or 37°C. After 48 hours of incubation, images of cells were taken and cell body diameter was measured. Cultures of each strain grown in YPD medium at 24°C examined at exponential phase of growth served as controls (A) Incubation at both 30°C and 37°C led to a significant difference in cell body diameter for both strains as compared to the control treatment (p<0.001). Incubation at 37°C led to a significantly larger median cell body diameter of H99 strain as compared to 30°C (p<0.001). While incubation of *C. gattii* at 37°C led to the presence of a small population of particularly large cells that were absent after incubation at 30°C, the effect of higher temperature on the median diameter was not statistically significant. The median cell diameter of R256 cells at respective temperatures was significantly larger as compared to H99 strain at 30°C (p < 0.001) and 37°C (p < 0.01). (B) DIC images of cells treated for 48 hours at 37°C illustrating larger cell size of R256 as compared to H99 strain. (C) A schematic illustrating the stimulating effect of 37°C on titanisation induced by 10% FBS in the presence of 5% CO_2_. Bars represent 10 μm.

At 30°C, both *C. neoformans* var. *grubii* strain H99 and *C. gattii* strain R265 were capable of titanisation and the relative percentages of Titan cells and proliferation rates were similar to those at 37°C, confirming that host temperature is not essential for titanisation (Figures 1, 2, 3, Table S2 and S3). However, the average size of Titan cells was smaller for both strains at 30°C as compared to 37°C suggesting that the temperature of 37°C potentiates this process (Figure 1). Importantly, at 30°C none of the non-*neoformans* and non-*gattii* strains formed cells that could be classified as *bona fide* Titans according to previously established criteria (Figure 2, Table S3). Notably, none of these strains underwent significant enlargement (Figure 2). We observed morphological differences among the non-*neoformans* and non-*gattii* species under “titanisation” conditions (Figure 2). Most of *C. albidus* cells, uniquely among other species, became uniformly round and unbudded indicative of cell cycle arrest. *C. terrestris* and to a lesser extend also *C. aspenensis* were the only species that exhibited a large proportion of cells with a single enlarged vacuole. In four species (*C. laurentii*, *C. terreus*, *C. uniguttulatus*, and *S. cerevisiae*) there was a significant number of cells containing multiple vesicles reminiscent of autophagic bodies. For *C. curvatus* frequent pseudohyphae were observed mixed with elongated yeast cells (Figure 2). While *C. terreus* exhibited a highly heterogeneous morphology with some cells being enlarged, these enlarged cells unlike typical Titan cells, were often not round, lacked single large vacuole and often contained a large daughter cell (Figure 2). Such morphology suggested possible failure in polarity establishment and a delay in the final cell separation during mitosis. Together, these data suggest that *C. neoformans* and *C. gattii* are unique in their ability to form Titan cells when exposed to 10% FBS in PBS in the presence of 5% CO_2_.

**Figure 2.**
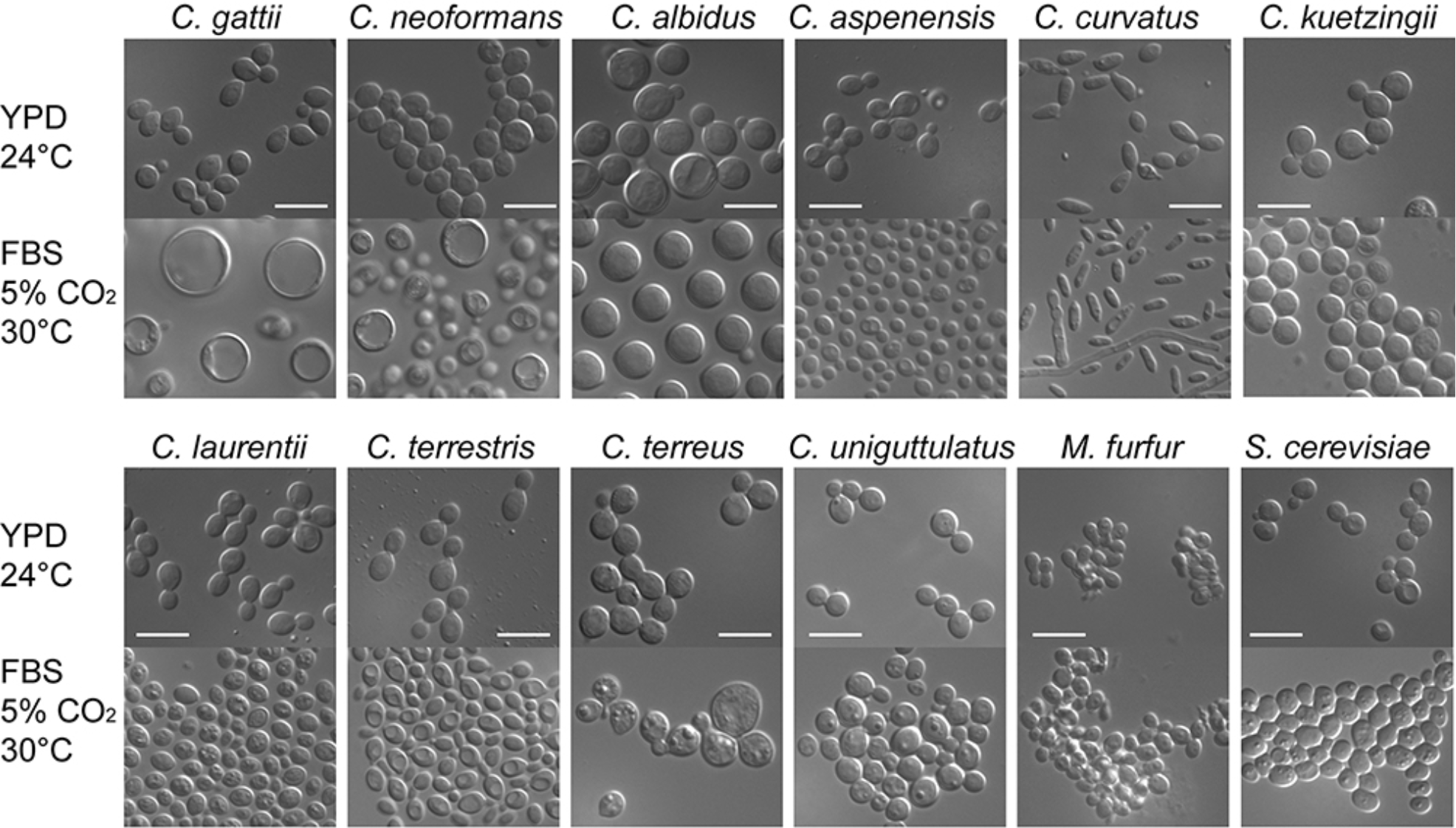
*C. neoformans* and *C. gattii* are unique in their ability to form Titan cells *in vitro* under the conditions involving 10% FBS in PBS, 5% CO_2_ at 30°C. *C. neoformans* var. *grubii* (H99), *C. gattii* (R265), and representatives of other species as indicated were subject to the *in vitro* titanisation protocol established by Dambuza et al. ^28^ with a modified temperature to 30°C. *C. neoformans* and *C. gattii* were the only two species that formed Titan-like cells. Bars represent 10 μm.

### FBS kills *C. neoformans* var. *grubii* and inhibits growth of *C. gattii* upon incubation at 37°C in the absence of 5% CO_2_

We noticed that when *C. neoformans* var. *grubii* was incubated under conditions similar to those established by Dambuza et al. ^28^ but without applying 5% CO_2_, nearly all cells died after 48 hours of incubation (Figure 3 and S1, Table S2). Killing was tested based on two criteria. First, while the initial culture density was ~10^3^ cells/ml, after 48 hours almost no cells were detected inside the inoculated micro-wells of the multiwell plate, as judged based on microscopic examination (Figure S1). This finding suggests that the majority of cells have lysed after 48 hours. Second, when after 48 hours the entire content of the micro-well was spread on the YPD semisolid medium, no more than 0.2% of the initial population developed into colonies after 3 days of incubation under standard growth conditions (Figure 3, Table S2). Killing was not instant as after 24 hours of incubation in FBS media, cells were still readily detected under the microscope, although growth was inhibited. As the medium contained only 10% FBS in PBS, this striking result suggested that FBS contains a factor (or several factors) that contributes to growth inhibition and killing. Notably, two *C. neoformans* var. *grubii* clinical isolates (H99 and MD31) and nine out of ten environmental isolates of this species (with the exception of the strain BS025, which was strongly inhibited but not killed) were killed by the FBS after 48 hours (Figure 3, Table S2). Our results were puzzling as Trevijano-Contador et al. also utilized FBS in their titanisation-inducing media and no killing was reported under conditions that did not include 5% CO_2_ ^27^. One potential explanation of this discrepancy was that the specific FBS we utilized was unique in being particularly inhibitory and possessing “killing” properties. To test this possibility, we compared four other FBS samples that were obtained from various manufacturers but represented the same type of active FBS. In addition, we included FBS that has been dialyzed to test if the “killing” potential was removed or reduced upon dialysis. Strikingly, FBS from all five sources but not the dialyzed FBS, killed two strains of *C. neoformans* var. *grubii* after 48h of incubation (Table 1). These findings suggest that the “killing” factor was present in various FBS samples and was removed in the process of dialysis.

**Figure 3.**
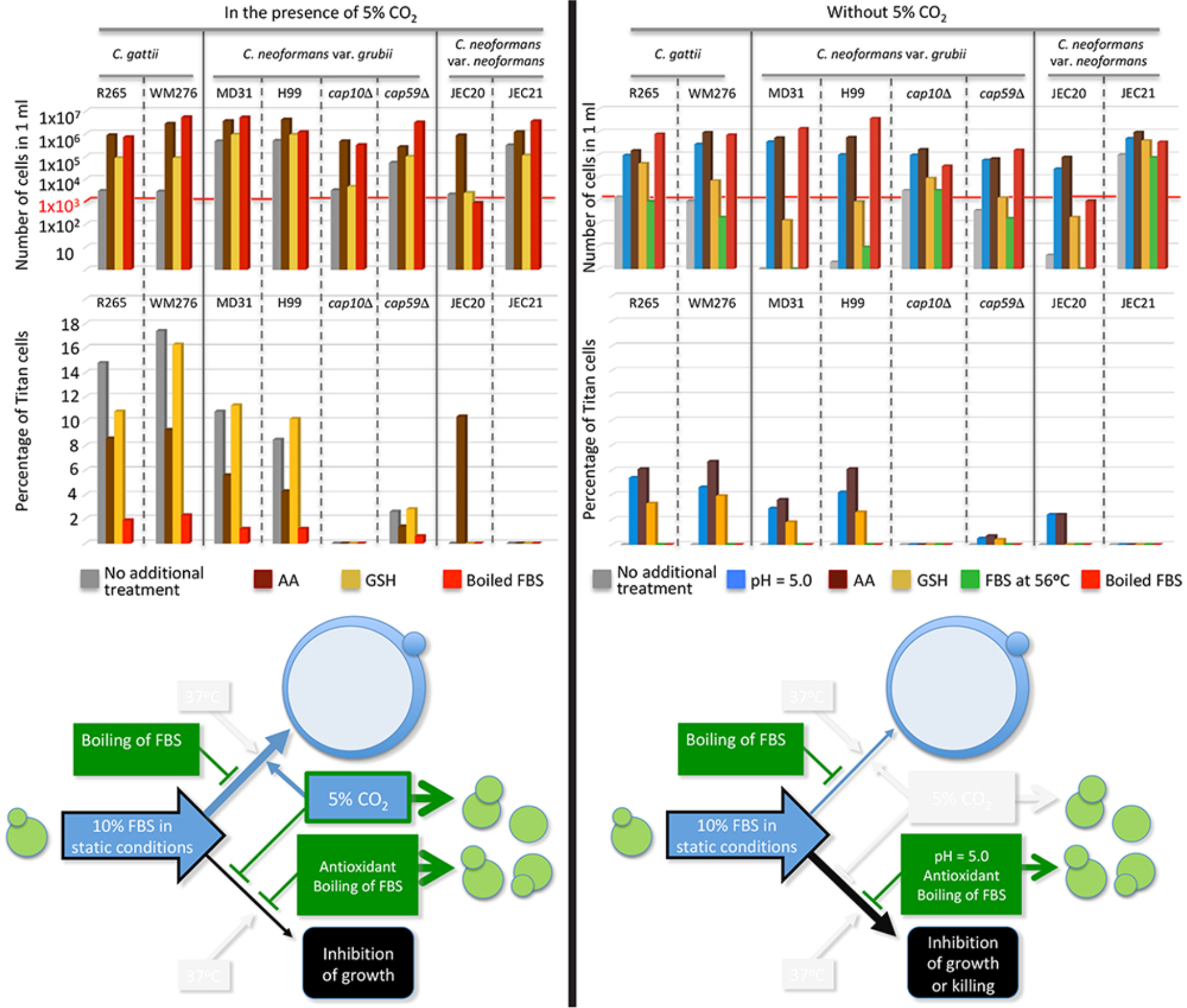
Fetal bovine serum possesses an inhibitory activity towards *C. neoformans* var. *grubii* and *C. gattii* and is sufficient to induce Titan cell formation in the absence of 5% CO_2_. Cells were subject to conditions established by Dambuza et al. either with or without the presence of 5% CO_2_ (indicated as “No treatment”). Titanisation experiment was also performed with the FBS or FBS-containing media subjected to the following additional treatments: FBS media were supplemented with either AA (10 mM ascorbic acid), GSH (5 mM reduced form of glutathione), or pH was adjusted to 5.0 (from the initial pH = 7.4); the FBS was brought to boiling point for 2 min or heated at 56°C for 30 minutes. Graphs are based on data included in Table S2, which contains more complete list of other strains tested. The schematic images illustrate proposed effects of the treatments based on findings presented in the corresponding graphs above each image. Green color in the schematic images refers to factors that generally reduce inhibition of growth imposed by the FBS. Bleu color refers to factors that stimulate titanisation. In the presence of 5% CO_2_ (shown on the left) Titan cells are formed (in strains that are capable of titanisation) and the inhibitory effect to the FBS is minimal. Boiling of FBS stimulates proliferation but inhibits titanisation whereas addition of an antioxidant reduces inhibition imposed by FBS without inhibiting titanisation. In the absence of 5% CO_2_ (shown on the right), the inhibitory effect of the FBS predominates and titanisation is inhibited unless pH is adjusted to 5.0, or an antioxidant is present. Boiling of the FBS reduces the inhibitory effect of FBS but also reduces titanisation.

**Table 1.** Cell viability (in the absence of 5% CO_2_) and percentage of Titan cells (in the presence of 5% CO_2_) of two *C. neoformans* var. *grubii* strains incubated with various types of sera (FBS) ^#^

| Tested strains | Fetal bovine serum (name of the company, catalog number) |  |  |  |  |  |
| --- | --- | --- | --- | --- | --- | --- |
|  | Atlas Biol.<br>F-0500-A | Atlanta Biol.<br>S11150 | Corning<br>35-075-CV | VWR<br>89510-186 | Sigma<br>F6178 | Gibco*<br>26400-044 |
|  | Viability (percent survived) after 48 h incubation at 37°C without 5% CO<sub>2</sub> |  |  |  |  |  |
| H99 | 0% | 0% | 0% | 0% | 0% | 73.8% |
| Btc012Gbc98-1 | 0.5% | 0.4% | 0% | 1.1% | 1.4% | 76.5% |
| Percentage of Titan cells after 48 h incubation at 37°C with 5% CO<sub>2</sub> |  |  |  |  |  |  |
| H99 | 8.6% | 9.4% | 9.6% | 8.7% | 7.3% | 0.4% |
| Btc012Gbc98-1 | 7.9% | 9.7% | 8.2% | 5.9% | 6.8% | 0.3% |

<sup>#</sup>Tests were performed according to the protocol by Dambuza et al., except for cell viability measurements where 5% CO<sub>2</sub> has not been applied <sup>28</sup>.
\* Dialyzed FBS, which has no glucose and amino acids

While *C. gattii* was not killed by the FBS, growth of cells was strongly inhibited and no Titan cells were observed in the absence of 5% CO_2_ (Figure 3, Tables S2, S3). A lack of titanisation in case of *C. gattii* could be explained by a strong inhibitory effect of the FBS. However, when the boiled FBS was utilized, significant proliferation was observed, but no Titans were formed (Figure 3, Table S2). This suggests that FBS is not sufficient to induce titanisation in the absence of CO_2_ or alternatively, boiling not only eliminated inhibitory potential but also a factor (or factors) that stimulates titanisation. The latter could occur if the same factor (or factors) caused both, growth inhibition/killing and titanisation, but other alternatives remained possible. The two strains of *C. neoformans* var. *neoformans* showed a distinct response to FBS in the absence of 5% CO_2_; strain JEC21 was not inhibited while strain JEC20 was killed by FBS (Figure 3, Table S2). Importantly, strain JEC21 did not form Titan cells even though proliferation has occurred (Figures 3, S2 and Table S2).

We tested how does reducing the temperature to 30°C impact the inhibitory effect of the FBS. The two *C. neoformans* var. *grubii* strains tested under these conditions (H99 and MD31) were less affected by the FBS as 30-40% of the initial population of cells survived after 48 hours compared to nearly complete killing at 37°C (Table S3). *C. neoformans* var. *neoformans* strain JEC21 was not affected similarly to 37°C, whereas strain JEC20 remained largely inviable (Figure S2, Table S3). The two *C. gattii* strains remained strongly inhibited. Notably, FBS had no significant inhibitory effect on other non-*neoformans Cryptococcus* species, except for *C. uniguttulatus* strain MD26 and *C. terreus* (Table S3). FBS did not affect growth of *S. cerevisiae*, whereas proliferation of *M. furfur* and *M. sympodialis* was inhibited (Table S3). Our data suggest that FBS contains a factor or factors that kill *C. neoformans* var. *grubii* and strongly inhibit proliferation of *C. gattii*, *C. uniguttulatus*, *C. terreus*, *M. furfur*, and *M. sympodialis* at 37°C in the absence of 5% CO_2_. Incubation under 5% CO_2_ improves proliferation of the majority of strains tested, except for *C. uniguttulatus* strain MD26 and *C. terreus* (Tables S2 and S3).

### FBS can induce Titan cell formation in the absence of CO_2_

Previous studies have demonstrated that FBS contains components that trigger titanisation ^27,28,34^. Are the same components also responsible for growth inhibition and/or killing in the absence of 5% CO_2_? If the two activities had a separate origin, then hypothetically, eliminating the inhibitory effect of the FBS without affecting the factor (or factors) that induces titanisation should allow formation of Titan cells in our assay even in the absence of 5% CO_2_. We selected one of the FBS samples to test these possibilities. The results for two representative strains from each *C. neoformans* (serotype A and D) and *C. gattii* are included in Figure 3, whereas the results for the remaining strains that were tested are included in Table S2. We first considered that an enzymatic activity may be involved and we subjected the FBS to a brief boiling prior to utilizing it in our protocol. As mentioned before for the *C. gattii* strains, this treatment abolished growth inhibition of all strains tested but no Titan cells were formed suggesting that this FBS treatment abolished both activities (Figure 3, Table S2). All of the FBS samples we utilized are active (not heat inactivated). Therefore, we tested if inactivating the complement complex by heating the FBS at 56°C for 30 minutes ^35^ would eliminate killing/inhibition of growth without affecting titanisation, however this treatment had no effect (Figure 3, Table S2). It remained still possible that the heat inactivation process performed by the manufacturer is more effective. Therefore, we tested commercially available heat inactivated FBS for its potential to kill two *C. neoformans* var. *grubii* strains, H99 and Btc012. Interestingly this heat-inactivated FBS exhibited less inhibitory effect as ~52% of H99 and ~89% of Btc012 cells survived after 48 hours when this FBS was utilized in contrast to 0.2% of H99 and 15% of Btc012 survival rate when the active FBS was added to the media (Figure S3). This result suggests that while heat inactivated FBS remains highly inhibitory towards growth of *C. neoformans* var. *grubii* in the absence of 5% CO_2_, heat inactivation reduces its “killing” potential.

We further utilized the H99 and Btc012 strains to test how the initial cell density and the nature of the pre-incubation medium influence the inhibitory effects of the active FBS. To test the effect of the initial cell density, we inoculated either 10^3^ or 10^5^ cells into the micro-well and assessed cell viability after 48 hours of incubation with 10% FBS in the absence of 5% CO_2_. Strikingly, cells that were inoculated at 10^5^ were no longer inhibited by the FBS, suggesting that initial low cell density is critical for the killing effect to take place (Figure S3). We tested if the amount of amino acids in the YNB that was used as the pre-incubation medium (YNB with vs. YNB without amino acids) played a role in the subsequent inhibition by the FBS but found no significant difference (Figure S3). In contrast, utilizing rich YPD medium instead of YNB as the pre-incubation medium completely eliminated the inhibition (Figure S3). Importantly, under both of the conditions that allowed robust proliferation (inoculating 10^5^ cells or utilizing YPD as the pre-incubation medium) no Titan cells were detected in the absence of 5% CO_2_.

We hypothesized that growth inhibition and/or killing resulted from an increase in levels of reactive oxygen species (ROS). To test this possibility we first included in our “titanisation” media the reduced form of glutathione (GSH) as an antioxidant ^36^. Consistent with the role of ROS in growth inhibition caused by the FBS, addition of GSH reduced inhibition of growth in *C. gattii* and eliminated killing and allowed proliferation of *C. neoformans* var. *grubii* (Figure 3, Table S2). Importantly, addition of GSH allowed Titan cell formation in both *C. gattii* strains, and most of the *C. neoformans* var. *grubii* strains, although the percentage of Titans was always significantly lower as compared to conditions that included 5% CO_2_ (Figure 3, Table S2). Two environmental *C. neoformans* var. *grubii* strains (BS001 and BS042) showed significantly higher growth improvement in the presence of GSH (~100 times higher density compared to other strains). Strain BS001 did not form Titans under these conditions confirming that this strain is not capable of titanisation. In contrast, BS042 formed ~3.2% Titan cells despite relatively high density (Table S2). Interestingly, while addition of GSH further improved growth of *C. neoformans* var. *neoformans* strain JEC21, titanisation in this strain was not observed (Figures 3, S2, Table S2). Strain JEC20 remained inhibited, suggesting that some factor other than ROS contributes to growth inhibition in this strain (Figure 3, Table S2). To further test the role of ROS we utilized ascorbic acid (AA) as an alternative antioxidant ^37^. Strikingly, all the strains that exhibited better growth and titanisation in the presence of GSH showed even more significant growth improvement and consistently a higher percentage of Titan cells upon addition of 10 mM AA (Figure 3, Table S2). Theoretically, AA may be a more potent antioxidant (either by reducing ROS or alleviating the negative effects of ROS) as compared to GSH at concentrations that were applied. One notable difference between the two treatments was that the addition of AA to FBS lowered the pH from ~7.4 to ~5.0, while GSH had no effect on pH. To test if the potentiated effect of AA was due to lower pH, we adjusted the pH of the FBS-containing media to 5.0 with hydrochloric acid (HCl). Strikingly, FBS-containing medium adjusted to pH 5.0 was no longer inhibitory and allowed titanisation of most of the strains (Figure 3, Table S2). Notably, the effect of lowering the pH was intermediate between the effect of GSH and AA for all except three *C. neoformans* var. *grubii* strains (BS244, BS042, BS051) for which the effect of AA was less pronounced (Figure 3, Table S2). Strikingly, either an addition of AA or lowering pH to 5.0 not only allowed proliferation of the strain JEC20 but also led to formation of Titan cells, suggesting that JEC20 is significantly more sensitive to the inhibitory effects of FBS but nonetheless capable of titanisation (Figures 3 and 4).

**Figure 4.**
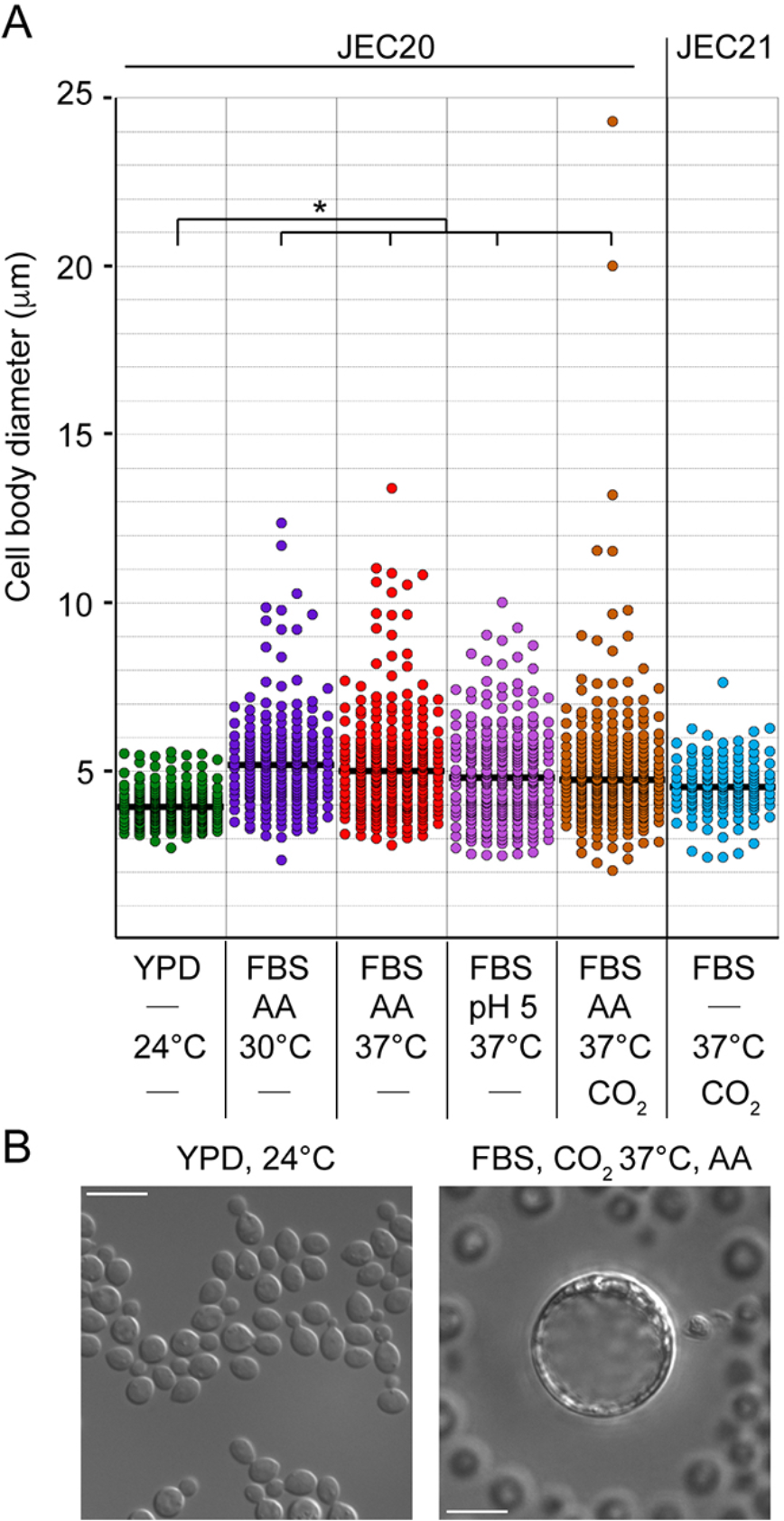
*C. neoformans* var. *neoformans* strain JEC20 is capable of titanisation under conditions with pH adjusted to 5.0. Cells were subject to the protocol by Dambuza et al. ^28^, except the FBS-containing medium was treated as indicated. Adjusting pH to 5.0 or addition of 10 mM ascorbic acid (AA) led to a significant increase of the median cell body diameter (p < 0.001). Addition of AA at 37°C under 5% CO_2_ led to appearance of particularly large Titan cells. In contrast, strain JEC21 was not able to form Titan cells. (B) Strain JEC20 forms Titan cells under modified conditions by Dambuza et al. by addition of 10 mM ascorbic acid (AA). Bar represents 10 μm.

We found it striking that *C. gattii* was not killed by the FBS in contrast to *C. neoformans* var. *grubii*. *C. gattii* has been shown to possess relatively higher resistance to oxidative stress due to specific mitochondrial morphology ^38^. Notably in the protocol proposed by Trevijano-Contador et al. sodium azide has been utilized ^27^. As sodium azide inhibits mitochondrial function and may reduce levels of ROS, this may explain why Trevijano-Contador et al. have not observed growth inhibition by FBS in the absence of 5% CO_2_. To test this possibility, we added sodium azide to our titanisation medium and evaluated growth inhibition and killing after 48 hours. Sodium azide at a concentration equal to 15 μM did not rescue growth in the absence of 5% CO_2_ (Table S4). However, it improved proliferation in the presence of 5% CO_2_ suggesting that mitochondrial activity contributes to growth inhibition in the presence of FBS (Table S4).

As shown in Table 1, dialyzed medium was not killing *C. neoformans* var. *grubii* in the absence of 5% CO_2_, in contrast to non-dialyzed types of FBS, suggesting that the factor (or factors) that contribute to killing was removed during dialysis. However, no proliferation was observed most likely due to the absence of glucose and amino acids that were removed during the dialysis procedure. Indeed, when the dialyzed medium was supplemented with glucose and amino acids, proliferation of C. *neoformans* var. *grubii* was evident (Table S4). This dialyzed FBS that was supplemented with glucose and amino acids fully supported titanisation in the presence of 5% CO_2_. However, when this FBS was subject to boiling the percentage of Titan cells was significantly lowered (Table S4). This finding suggests that this dialyzed FBS still contained a factor (or factors) that stimulate titanisation in the presence of 5% CO_2_. On the other hand, in the absence of 5% CO_2_, no Titans were formed with this dialyzed FBS despite robust proliferation, which indicates that dialysis eliminated another factor that normally is present in FBS and which is sufficient to stimulate titanisation in the absence of 5% CO_2_ (Table S4).

Previous studies have suggested that the polysaccharide capsule plays an important role in Titan cell formation, potentially serving as a receptor for the external signal that triggers titanisation ^32^. We hypothesized that capsule provides also a receptor for the signal that causes growth inhibition imposed by the FBS. To test this possibility we subjected two acapsular mutants, *cap10*Δ and *cap59*Δ to the *in vitro* titanisation conditions. Consistent with previous studies, under titanisation conditions, *cap59*Δ failed to form capsules, whereas *cap10*Δ formed capsules that were significantly reduced as compared to the capsule size of the congenic H99 wild type strain (Figure 5). Strikingly, both mutants were partially resistant to growth inhibition imposed by the FBS (Figure 3). Moreover, *cap59*Δ mutant exhibited reduced ability to titanize while *cap10*Δ was not able to produce Titans under any conditions tested (Figure 3 and Figure 5). In aggregate, our data suggest that *C. neoformans* and *C. gattii* are unique in being able to titanize in the presence of the FBS and for their hypersensitivity to FBS. FBS possesses an activity that is capable of stimulating titanisation even in the absence of CO_2_ and which may be the same activity that causes growth inhibition (Figure 6). The inhibitory and killing effects of FBS result in part from an increase in intracellular ROS and are dependent on pH and temperature. Specifically, growth inhibition is more potent at neutral pH at 37°C as compared to acidic pH and 30°C. The growth inhibition or killing effect caused by FBS in the absence of 5% CO_2_ prevents titanisation unless cell proliferation is rescued by lowering pH or addition of an antioxidant. Furthermore, growth in the presence of 5% CO_2_ reduces the inhibitory effect of FBS and further stimulates titanisation (Figure 6).

**Figure 5.**
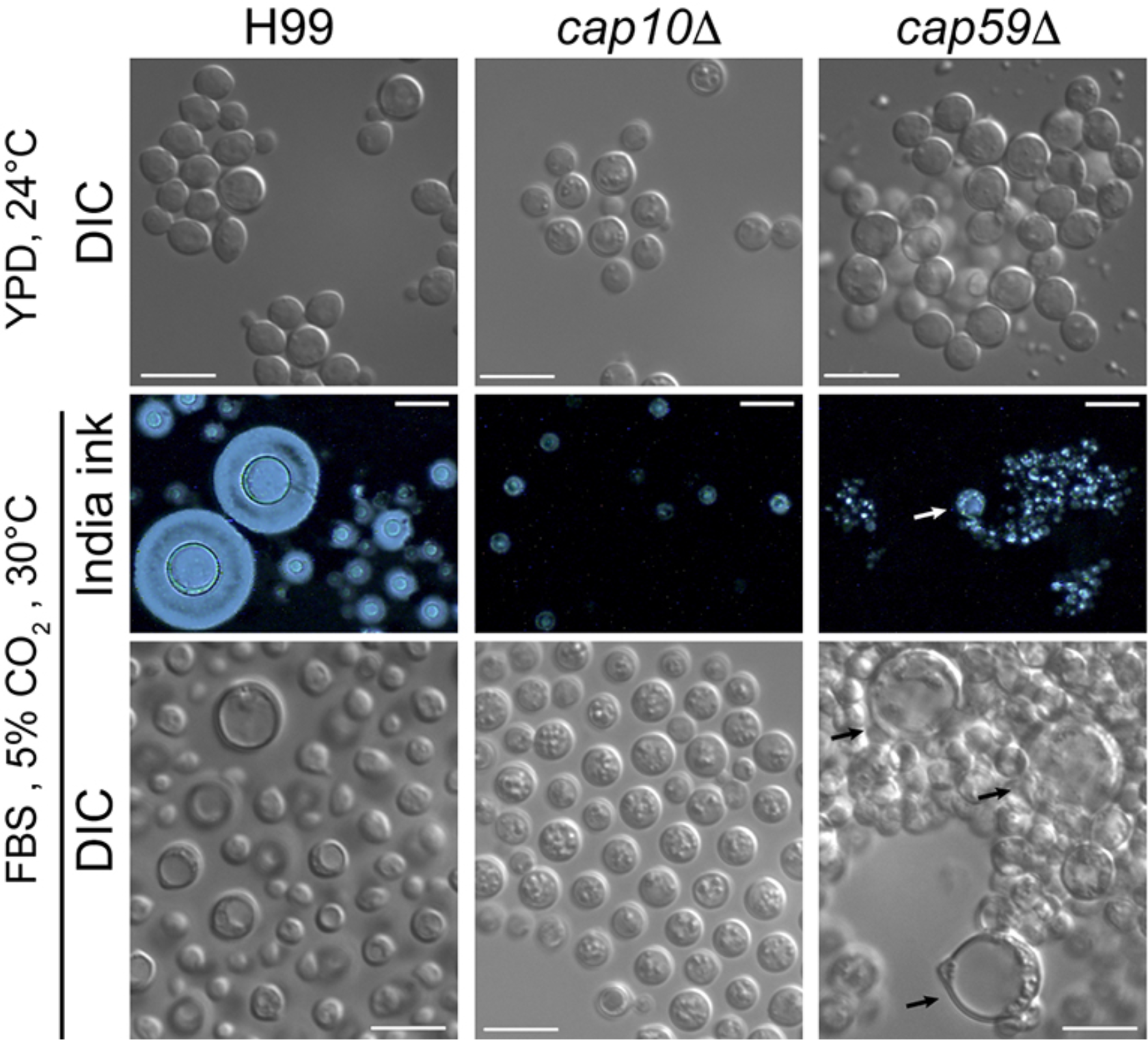
*C. neoformans* var.*grubii* acapsular mutants *cap10*Δ and *cap5*Δ exhibit distinct responses to titanisation media under standard conditions. Wild type strain (H99) and the two mutants were incubated in YPD at 24°C (control) or in titanisation media ^28^ at 30°C under 5% CO_2_ for 48 hours. Under control conditions *cap59*Δ cells formed aggregates whereas H99 and *cap10*Δ cells did not. India ink staining indicated a complete lack of capsule in *cap59*Δ and significantly reduced capsule in *cap10*Δ in agreement with previous findings ^28^. Incubation in titanisation media led to formation of Titan cells in H99 and the *cap59*Δ mutant but not in the *cap10*Δ mutant. Bars represent 10 μm.

**Figure 6.**
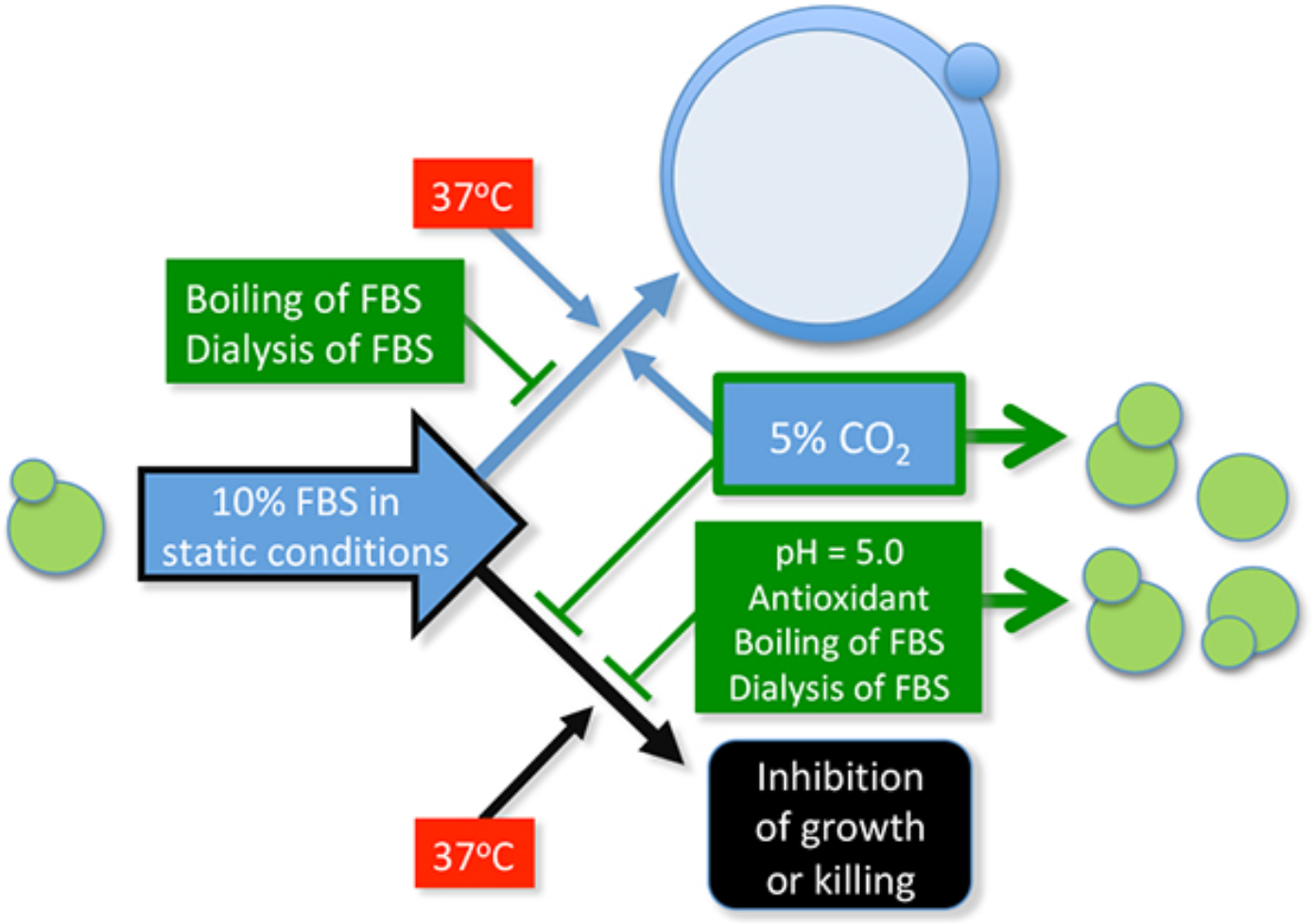
A schematic illustration of a unique response of *C. neoformans* and *C. gattii* to *in vitro* titanisation conditions involving FBS. Green color refers to factors that reduce the inhibitory effect of FBS and promote proliferation. Blue color relates to factors that stimulate titanisation. Under static growth conditions, 10% FBS (in PBS) possesses an activity that is capable of stimulating titanisation even in the absence of CO_2_. However, an inhibitory activity of FBS precludes titanisation unless this inhibitory effect is diminished by at least one of the following treatments: addition of an antioxidant, lowering pH to 5.0, or introduction of 5% CO_2_. Furthermore, 5% CO_2_ not only reduces the inhibitory effect of FBS but also further stimulates titanisation. Boiling of the FBS eliminates both, the inhibitory effect of the FBS and the activity that induces titanisation with and without the presence of 5% CO_2_. Dialysis of the FBS also prevents inhibition imposed by the FBS and prevents titanisation in the absence of 5% CO_2_. However, dialyzed FBS still allows titanisation in the presence of 5% CO_2_. A temperature of 37°C has the opposite effect as it potentiates both, titanisation and the inhibitory effect of the FBS. Capsule plays both a positive role by enhancing titanisation and also a negative role by mediating the inhibitory effects of the FBS (not shown).

## DISCUSSION

The leading aim of this study was to assess the ability to titanize among various non-*C. neoformans*/non-*C. gattii* yeast species. We utilized the *in vitro* titanisation protocol described recently by Dambuza et al. ^28^. Knowing that *C. neoformans* and *C. gattii* are unique among species within the order Tremellales for their ability to grow at temperature above 30°C ^39^ and based on our tests (Table 1) it was necessary to slightly modify this protocol by changing the incubation temperature from 37°C to 30°C. We demonstrated that eleven out of twelve of the tested strains of *C. neoformans* var. *grubii* and *C. gattii* strains (including reference strains) were able to form Titan cells at both 30 and 37°C.

Our results clearly show that under *in vitro* conditions examined here neither of evaluated strains of non-*C. neoformans*/non-*C. gattii* species is able to form Titan cells (Figure 2; Table S2, S3) as defined by adopted criteria ^28^. Moreover, the same experiment performed for comparison at 37°C confirmed that even strains capable of growing at this temperature (all the *C. laurentii* tested strains) were not able to form Titan cells. Changes at the micromorphology level were observed only in case of both tested strains of *C. curvatus*. While cultures incubated in YPD medium consisted of only single, elongated budding cells, in 10% FBS branched pseudohyphae similar to species belonging to *Trichosporon* genus were observed (Figure 2). It should be underlined that for our study we selected strains which represent species more or less phylogenetically related to the genus *Filobasidiella Kwon-Chung* (1976) which included two of the most important species (in the context of prevalence of cryptococcosis): *C. neoformans* and *C. gattii* ^7^. According to recently updated taxonomy of Tremellomycetes ^40^, strains used in this study belong to the following genera: Cutaneotrichosporon (*C. curvatus*), Filobasidiella (*C. gattii* and *C. neoformans*), Filobasidium (*C. uniguttulatus*), Naganishia (*C. albidus* and *C. kuetzingii*), Solicoccozyma (*C. terreus*), Papiliotrema (*C. aspenensis, C. laurentii, C. terrestris*). Among other basidiomycetous yeasts we also tested two reference strains that belong to *Malassezia* genus (*M. furfur* CBS 14141, *M. sympodialis* ATCC 42132). The *S. cerevisiae* BY4741 reference strain was used as outgroup (Table S1). While the strains tested here represent various groups within basidiomycetous yeasts, it is still possible that other species may be capable of titanisation. For instance more closely related *Cryptococcus* species *sensu stricto*, *C. amylolentus* may be able to titanize. Unfortunately, only one isolate of this species is available, which undermines the significance of this potential test. Additionally, it is plausible that some or all of the species tested in this study are capable of morphogenetic transition to form Titan cells but they may require specific environmental cue not tested here. While this alternative remains a possibility, we would like to note that formation a true hyphae, another morphological transition, is not common to all yeasts. While *C. albicans* forms enlarged “Goliath” cells in response to zinc limitation, this morphological transition is not conserved in *C. parapsilosis*, *C. lusitaniae*, and *Debaryomyces hansenii* and Goliath cells are not morphologically similar to Titan cells ^41^. Furthermore, the uniqueness of *C. neoformans* and *C. gattii* for titanisation under specific conditions described here emphasizes the success of these species to evolve as human pathogens. Future studies should reveal if other environmental cues exist to induce various yeast species to form Titan cells possessing all the characteristics described for *Cryptococcus* species complex. Several genes were recently implicated in titanisation of *C. neoformans* var. *grubii* ^27–29^. It would be of interest to establish if these genes are conserved in species tested here and other species outside of the *Cryptococcus* species complex.

Three inconsistencies apparent between the three recent *in vitro* studies were resolved in this study. First, Dambuza et al. reported a lack of titanisation in the *C. gattii* strain R265 suggesting that perhaps under their specific conditions *C. gattii* is not stimulated to form Titan cells ^28^. This was not the case in our study as two *C. gattii* strains (including R265) formed Titan cells under conditions similar to those utilized by Dambuza et al. Second, while Trevijano-Contador et al. indicated that *cap59*Δ mutant fails to form Titan cells under their conditions, Dambuza et al. observed Titan cells under conditions established in their laboratory ^27,28^. We confirmed that *cap59*Δ is capable of forming Titan cells under the latter conditions and similarly to Dambuza et al. we also observe that titanisation in this mutant is not as efficient as in the congenic H99 strain. Interestingly, we observed that in contrast to *cap59*Δ strain, another acapsular mutant, *cap10*Δ, is unable to form Titan cells. This suggests that the role of capsule in titanisation is a complex matter, potentially reflecting differences in how the *cap59*Δ and the *cap10*Δ mutant interact with human dendritic cells ^42^. Future studies should explain the differential response of these two acapsular mutants to titanisation conditions. Third, while FBS was a stimulant with respect to titanisation in the protocols established by Dambuza et al. and Trevijano-Contador et al., in the work by Hommel et al. FBS had an inhibitory effect towards Titan cell formation ^29^. We believe this discrepancy can be explained by the results presented here. Specifically, the protocol by Hommel et al. did not include 5% CO_2_, which might have contributed to the inhibitory effect of the FBS. Given that killing in our case was observed after ~48 hours, while Hommel et al. read their results after a few hours, the killing effect might have not been observed.

We believe our results provide a robust support for the inhibitory effect of the FBS. First, the effect was confirmed with the use of five FBS samples obtained from various manufacturers. Second, the effect was common to several independent strains of *C. gattii* and *C. neoformans* var. *grubii*. Third, an identical sample when incubated in the presence of 5% CO_2_ was no longer inhibited by the FBS. We provided strong evidence that the inhibition is resulting from FBS, as boiled or dialyzed FBS was no longer inhibitory. This brings an important question of why this inhibitory effect of FBS has not been reported by other laboratories. Clearly numerous studies have been utilizing FBS for incubation of *C. neoformans* under various conditions. We hypothesize that such studies did not report the inhibition or killing for the following possible reasons: 1. Many of such studies cultured the cells by shaking while our conditions are static. 2. Other studies incubated the cells under 5% CO_2_ while we observe killing in the absence of 5% CO_2_. 3. The results of other studies are often read after no longer than 24 hours, whereas killing in our case was detected at 48 hours. 4. Often, the heat-inactivated FBS is utilized, whereas we employed active FBS and our results suggest that heat inactivation reduces the inhibitory effect. 5. Other studies may set the yeast cultures at a relatively higher density; we established that at higher densities the inhibitory effect diminishes. Thus, potentially for the killing/inhibition to take place specific conditions need to be met: static growth (no shaking), 37°C, absence of 5% CO_2_, neutral or slightly alkaline pH, very low cell density (~10^3^ cells/ml), presence of active FBS. Interestingly, most of these factors are also critical to induce titanisation by the FBS as described recently ^27–29^.

Our results strongly suggest that *C. neoformans* and *C. gattii* are unique among other basidiomycetous yeasts for their ability to form Titan cells under specific conditions applied here 28. Strikingly, these two pathogenic species are also unique for their high susceptibility toward active FBS when cells are cultivated without the presence of 5% CO_2_. Furthermore, there are notable differences between the two species. Whereas almost 100% of *C. neoformans* var. *grubii* cells are killed after 48 hours of incubation in 10% FBS (w/o CO_2_), *C. gattii* cells remain still viable (confirmed by subsequent plating on YPD semisolid media) although they are strongly inhibited and not able to proliferate under such conditions. An exception to this rule was the response of two strains of *C. neoformans* var. *neoformans* (JEC20 and JEC21). JEC21 was resistant to the inhibition imposed by FBS and it was not able to titanize even in the presence of 5% CO_2_ and that way it resembled strains of non-*neoformans*/non-*gattii* species. Although under some conditions enlarged cells of JEC21 were found, these cells were distinct form *bona fide* Titans (Figure S2). JEC20 on the other hand was particularly sensitive to slightly alkaline pH and when the pH was established at 5.0, JEC20 proliferated and was capable of titanisation. This result points to a distinct nature of the two strains and further supports the relationship between sensitivity to FBS and the ability to titanize.

An important question is whether the inhibitory effects of the FBS and the activity stimulating titanisation have the same origin. We were able to reconstitute conditions where cells were no longer inhibited by the FBS in the absence of 5% CO_2_ and yet they still formed Titans, which may suggest that the two factors are separate. However, those conditions include addition of an antioxidant or lowering the pH, modifications that might have simply alleviated the inhibitory effect rather than eliminating the inhibitory activity per se. To further address this question, we examined the consequences of three ways of FBS modification: brief boiling, heat inactivation at 56°C, and dialysis. Boiled FBS lost the “killing” activity, while the activity to induce Titan cells was completely lost under conditions without 5% CO_2_ and significantly reduced with 5% CO_2_ (Figure 3, Table S2, S3). This is consistent with the same activity stimulating titanisation and inhibiting growth. While heat inactivation at 56°C when performed in the lab had no effect, FBS that was commercially heat-inactivated was less inhibitory suggesting that the complement complex may contribute to the killing and/or inhibition (Figure S3). Dialyzed serum reconstituted with glucose and amino acids was no longer inhibitory. According to the manufacturer’s specification data sheet all components of FBS with the size below 10,000 Daltons are eliminated during the dialysis process. This suggests that the molecule causing inhibition is not a protein. However, it is also possible that dialysis eliminates a component that is essential for inhibition by acting as a co-factor for certain enzyme. Interestingly, while the FBS that has been dialyzed did not stimulate titanisation in the absence of 5% CO_2_, it supported titanisation in the presence of 5% CO_2_. This finding suggests that FBS contains at least two factors, one that is sufficient to stimulate titanisation in the absence of 5% CO_2_ and which is lost upon dialysis and the other factor that is capable of stimulating titanisation in the presence of 5% CO_2_ and which is not lost upon dialysis. As dialysis eliminates molecules of molecular weight below ~ 10KD, this suggests that the first factor is not a protein. Therefore, the lost component upon dialysis may be a phospholipid or peptidoglycan as suggested recently ^27,28^. Hommel et al. have reported that addition of 5% FBS or L-a-phosphatidylcholine inhibited Titan cell formation in hypoxia-based titanisation conditions. Perhaps polar lipids present in FBS are responsible for both, titanisation and growth inhibition in the absence of 5% CO_2_. Trevijano-Contador et al. have proposed that phospholipids stimulate titanisation through a degradation product, diacylglycerol (DG), and subsequent activation of the PKC pathway by DG ^27^. Therefore, consistent with our results, phospholipids may be one of the FBS components that are sufficient to stimulate titanisation in the absence of 5% CO_2_ and which is lost upon dialysis. Furthermore, Dambuza et al. provides compelling evidence that bacterially derived peptidoglycan is sufficient to stimulate titanisation: exposure of cells to NMAiGn, a synthetic version of the bacterial cell wall component MDP, was sufficient to induce Titan cells, which would be also consistent with the results presented here ^28^. The same study presented evidence that the bronchoalveolar lavage fluid (BAL) also stimulated titanisation in place of the FBS. It would be of interest to test if BAL or bacterially derived cell wall components are inhibitory towards or kill *C. neoformans* under conditions of low cell density, 37°C, neutral pH, in the absence of 5% CO_2_.

If followed by the rule of parsimony, we should conclude based on former studies and the data presented here that the ability to form Titan cells and sensitivity to FBS are related and together constitute a unique feature of *Cryptococcus* species complex. Several important questions remain to be addressed by future investigations. First the identity of the factor present in FBS that inhibits growth of *C. gattii* and kills *C. neoformans* cells and the cellular pathways that mediate these responses are unknown. Second, it is presently unclear whether the factor that induces titanisation and the activity that inhibits growth have the same origin. As mentioned above, it is plausible that FBS contains more than one factor, one of which can stimulate titanisation in the absence of 5% CO_2_ and another activity that primes cells for titanisation in the presence of CO_2_. Third, it remains unclear what is the mechanism through which CO_2_ rescues growth *C. gattii* and *C. neoformans* in the presence of FBS. While we observed diminished growth inhibition upon setting the pH to 5.0, the effect of CO_2_ on pH may not account for the rescue. This is because the pH of the “titanisation” media under 5% CO_2_ was only slightly lower (changed from ~7.4 to ~ 7.2). We can speculate based on former studies that the rescue by CO_2_ is due to stimulation of the cAMP pathway, which is known to trigger cell wall remodeling in response to stress ^43^. This hypothesis is supported by our findings showing rescued proliferation in the presence of an antioxidant. Another aspect that needs more investigation is the multifaceted involvement of the capsule in titanisation. Our results of the experiments involving the acapsular mutants confirm established view that capsule plays an important role in titanisation. The novel aspect derived from our results is that the presence of capsule makes cells more sensitive to the inhibitory activity of FBS. Perhaps a specific capsule-dependent receptor or receptors govern these two distinct responses. Finally, our study prompts an important question of whether the unique sensitivity of *Cryptococcus* species complex towards FBS has any significant implications for human infection. Unusual sensitivity of *C. neoformans* var. *grubii* towards a component of the FBS potentially also present in human BAL or/and serum, given its success to cause cryptococcosis seems counterintuitive. Future studies should shed more light on the relatedness of the titanisation induction and the inhibitory effects of the FBS, the role of capsule in these cellular reactions, and the significance of these responses for cryptococcosis.

## MATERIALS AND METHODS

### Tested strains and media

All the strains used in this study are listed in Table S1, which includes information about the origin (source) and actual taxonomic position for each tested strain. Strains were routinely cultured on YPD semi-solid medium, followed by liquid medium: 1% yeast extract, 2% bacto-peptone and in case of semi-solid media 2% bacto-agar (BD Difco, Sparks, MD, USA, cat. no. REF212720, REF211820 and cat. no. REF212720, respectively) and filter-sterilized 2% glucose (VWR International LLC, West Chester, PA, USA, cat. no. BDH0230). In case of Titan induction experiment, cells were grown overnight in 5 ml YNB liquid medium: 0.67% Yeast Nitrogen Base with amino acids, pH 5.5 (Sigma cat. no. Y1250, St. Louis, MO, USA), 2% glucose at 30°C with horizontal shaking at 220 rpm (Thermo Scientific, MAXQ4450).

### *In vitro* Titan cells induction

Titan cells were generated *in vitro* according to recently described protocol ^28^. Experiments were performed in sterile 10% Fetal Bovine Serum (FBS, Sigma, cat. no. F6178) diluted in 1x concentrated PBS (Dulbecc’o Phosphate-Buffered Saline w/o Ca^2+^ and Mg^2+^, cat. no. REF21-031-CV, Corning cellgro®, Manassas, VA, USA), at final pH equal to 7.4. Original concentrated FBS was normally stored in 2.5 ml aliquots at −20°C to prevent repeated freeze-thaw procedure.

For comparison, to check the killing properties and efficiency of Titan cell induction several fetal bovine sera, from different companies were utilized (Table 1). Cells were incubated in static conditions for 48 h at 37 or 30°C and under 5% CO_2_ atmosphere (New Brunswick an Eppendorf company, Galaxy 170S, Ayrshire, Scotland) or w/o CO_2_ (Thermo Scientific, Heratherm Incubator IMH180, Langenselbold, Germany). If necessary described herein conditions were modified as follows. 1 M HCl (Sigma cat. no. 320331) was used to lower the pH to 5.0. To check the influence of antioxidants on viability and Titan cells formation serum was supplemented with the reduced glutathione (GSH, Fisher Scientific, cat no BP25211) and L-ascorbic acid (AA, Fisher Chemical, cat. no. A61-100) at a final concentration 5 mM and 10 mM, respectively. Sodium azide (NaN_3,_ Acros Organics, cat. no. 190380050, Belgium) was added to FBS at final concentration equal to 15 μM. Serum inactivation was performed in two ways. To obtain the boiled serum the concentrated serum was brought to boiling point and kept under such conditions for 1-2 min. Heat inactivated serum was obtained by heating the active serum in water bath (set to 56°C) for 30 minutes.

### Evaluation of cell viability

Viability of cultures was evaluated qualitatively by examining micro-wells from the titration plate directly under the microscope and quantitatively by plating. For this purpose, 100 μl volume from each micro-well of the titration plate was collected (after mixing) and either initially diluted if necessary using sterile 1x concentrated PBS or directly spread on YPD semi-solid medium. Plates were incubated 48 h at 30°C in the incubator w/o CO_2_ (Thermo Scientific, Heratherm Incubator IMH180, Langenselbold, Germany). After this time plates were photographed and colony counting was performed.

### Microscopy and imaging

Microscopic observations were performed using Zeiss microscope using either the 20x or 100x objective (Axiovert 200M, Zeiss ID#M 202086, Carl Zeiss MicroImaging, Inc., Thornwood, NY, USA) with built-in camera for photographic documentation (AxioCam HRm) and interfaced with AxioVision Rel 4.8 software (Carl Zeiss, Thornwood, NY). To visualize capsule a standard test with the use of india ink staining was performed and cells were photographed using dissection microscope Sporeplay (Singer Instruments, UK), For evaluation of cell body diameter cells were measured based on the images using ImageJ (https://imagej.nih.gov/ij/) and the data was visualized using the KaleidaGraph software (Synergy, Reading, PA).

### Statistical analysis

To test differences between the median of cell body diameter, Mann-Whitney test for non-parametric distributions was performed with the use of Kaleidagraph (Synergy, Reading, PA).

## ACKNOWLEDGMENTS

The authors thank Dr. Joseph Heitman for providing some of the non-*neoformans* species and Dr. John Perfect for providing environmental strains of *C. neoformans* var. *grubii*. The authors thank Dr. Sophie Altamirano, Dr. Elizabeth Ballou, and Dr. Kirsten Nielsen for helpful comments on the manuscript. RCR is partially supported by NIH grant 1R15 AI119801-01. LK is partially supported by NIH grants 1R15 AI119801-01 and 1P20GM109094-01A1.

## AUTHOR CONTRIBUTIONS STATEMENT

L.K. and M.D. designed the experiments. M.D performed most experiments, except those that are included in Figure S3 (performed by R.C-R). M.D. and L.K. wrote the main manuscript text and L.K prepared figures. All authors reviewed the manuscript.

## Supplementary Material

### Dylag M, Colon-Reyes R, Kozubowski L, Fetal bovine serum-triggered Titan cell formation and growth inhibition are unique to the *Cryptococcus* species complex

**Table S1.**
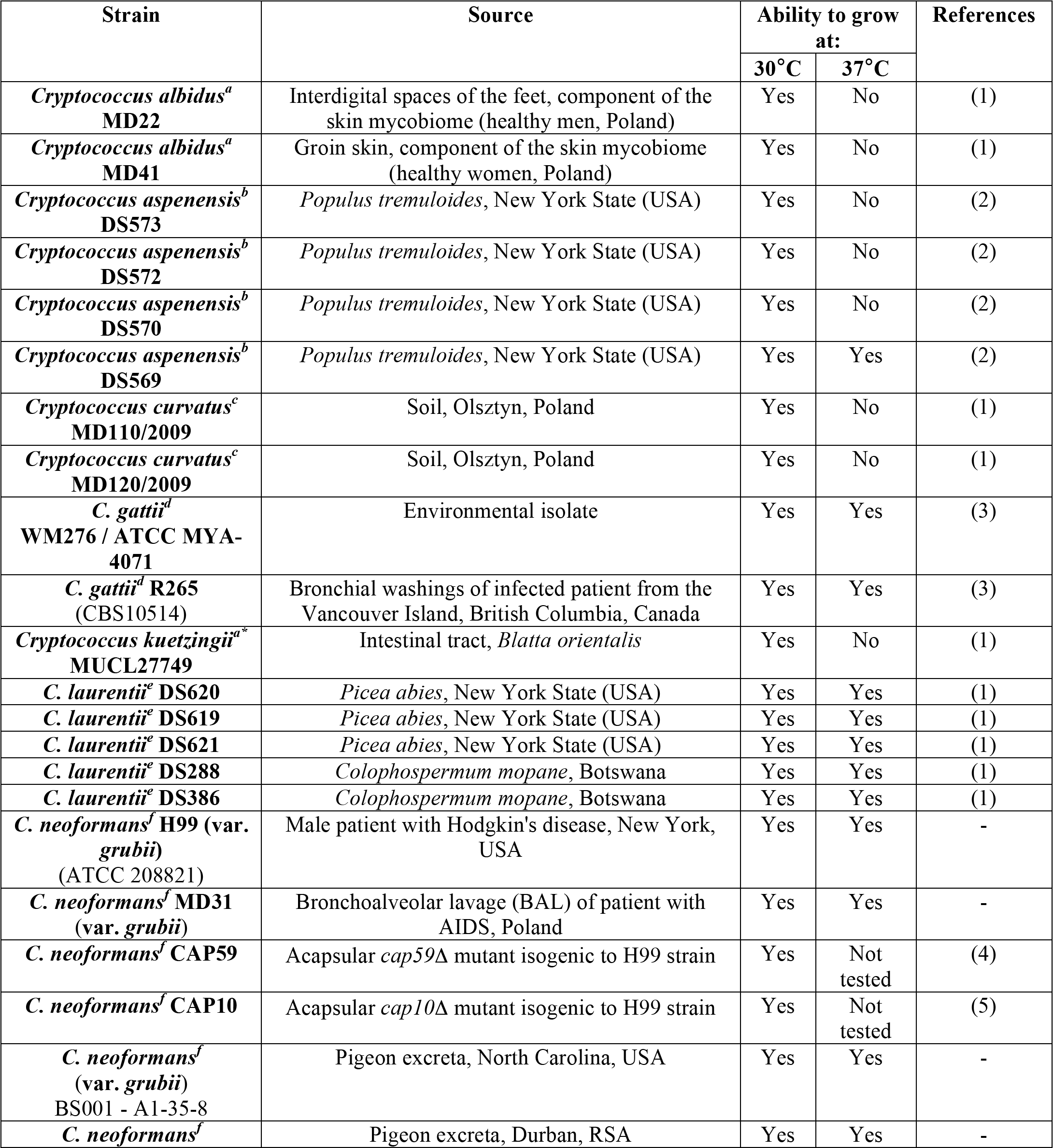
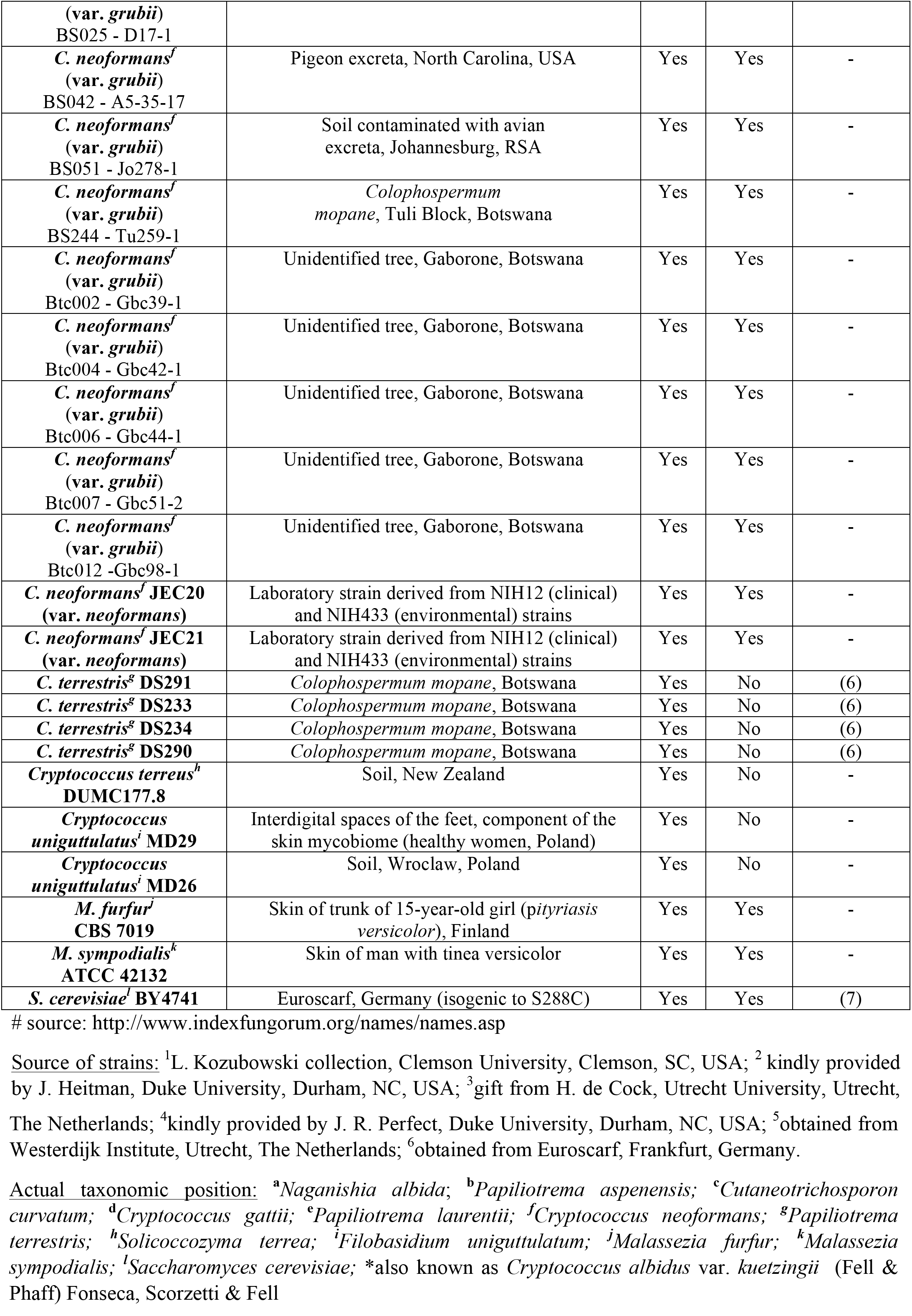
Strains used in this study - their source and actual taxonomic position^#^

**Table S2.**
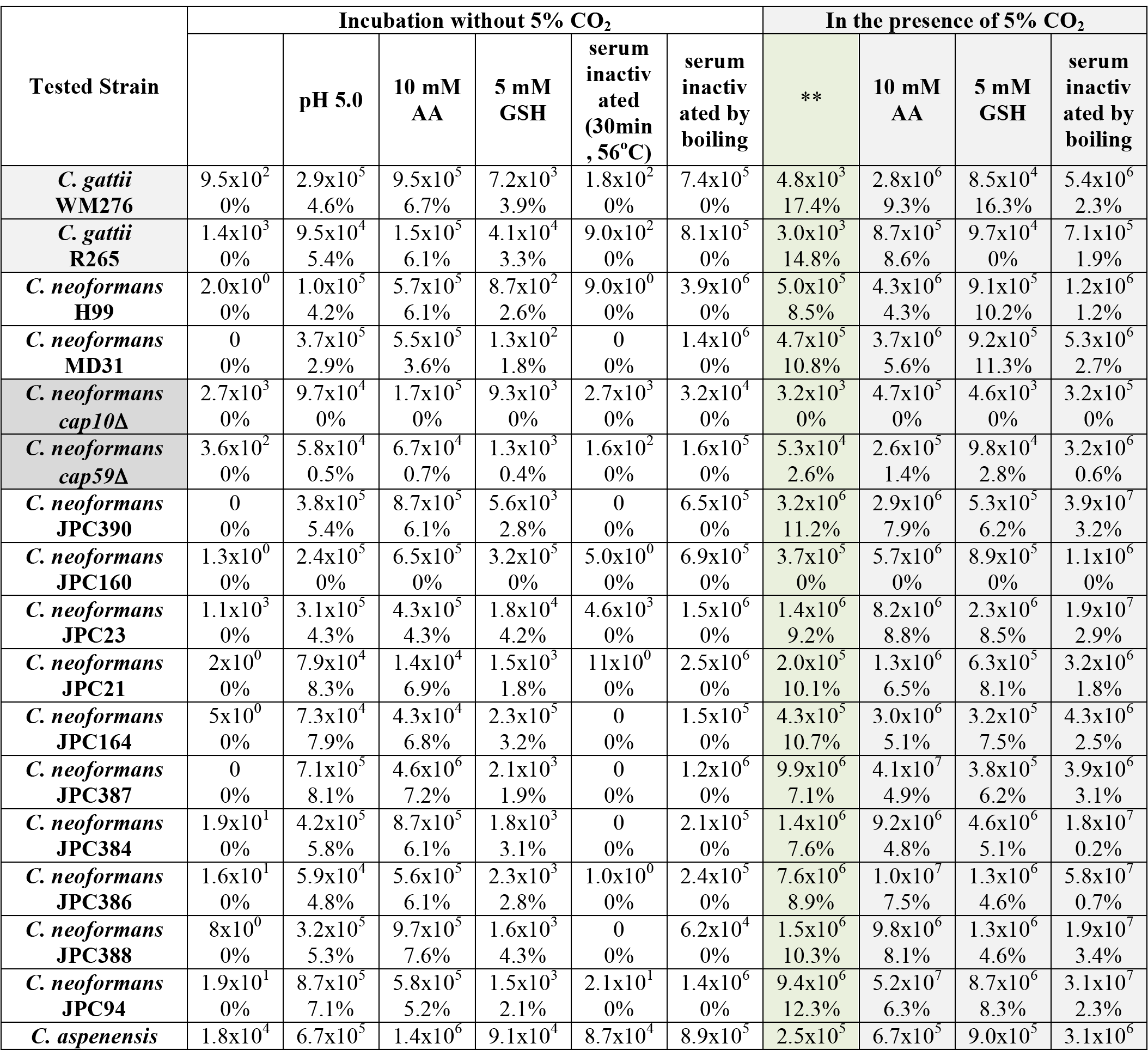
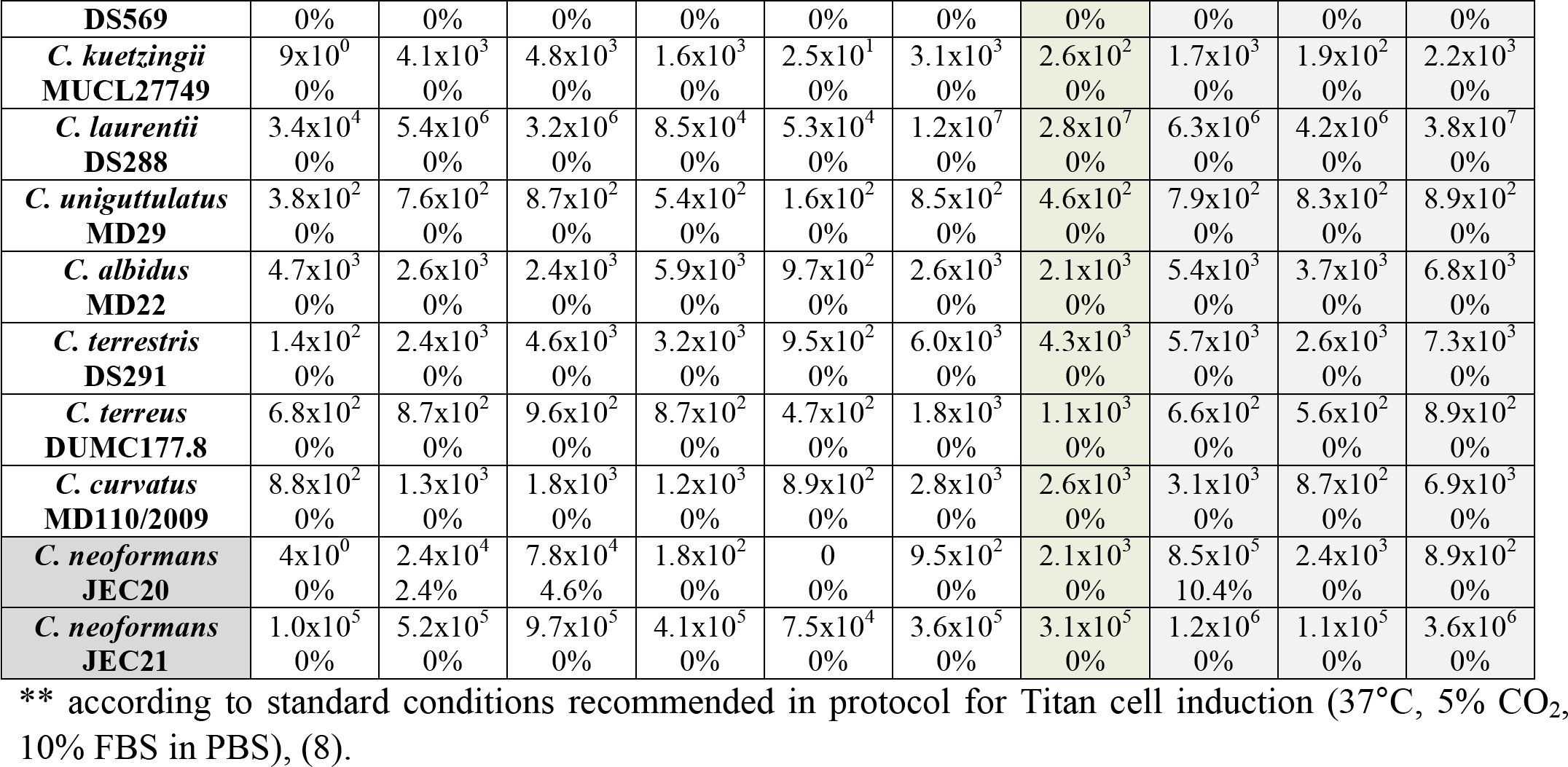
Total number of cells and percentage of Titan cells after 48 h incubation at 37°C under standard** and modified conditions. The initial inoculum was 10^3^ cells. Green-shaded column represents conditions similar to Dambuza et al. (8).

**Table S3.** Total number of cells and percentage of Titan cells after 48 h incubation at 30°C under various conditions. The initial inoculum was 10^3^ cells.

| Strain tested | Incubation without 5% CO<sub>2</sub> |  |  | In the presence of 5% CO<sub>2</sub> |  |  |
| --- | --- | --- | --- | --- | --- | --- |
|  |  | pH 5.0 | serum heat inactivated (30min, 56°C) | serum inactivated by boiling | ** | pH 5.0 |
| *C. gattii*<br>**WM276** | 1.1x10<sup>3</sup><br>0% | 1.5x10<sup>6</sup><br>5.1% | 9.7x10<sup>2</sup><br>0% | 1.1x10<sup>6</sup><br>0% | 1.5x10<sup>4</sup><br>18.5% | 4.3x10<sup>6</sup><br>16.3% |
| *C. gattii*<br>**R265** | 6.3x10<sup>3</sup><br>0% | 1.7x10<sup>5</sup><br>4.2% | 4.3x10<sup>3</sup><br>0% | 3.4x10<sup>5</sup><br>0% | 5.4x10<sup>3</sup><br>12.6% | 9.8x10<sup>5</sup><br>15.1% |
| *C. neoformans*<br>**H99** | 4.3x10<sup>2</sup><br>0% | 1.9x10<sup>4</sup><br>1.5% | 6.2x10<sup>2</sup><br>0% | 1.3x10<sup>6</sup><br>0% | 2.2x10<sup>5</sup><br>7.2% | 2.4x10<sup>6</sup><br>2.8% |
| *C. neoformans*<br>**MD31** | 3.1x10<sup>2</sup><br>0% | 6.6x10<sup>4</sup><br>1.5% | 5.2x10<sup>2</sup><br>0% | 4.3x10<sup>6</sup><br>0% | 1.2x10<sup>5</sup><br>8.4% | 1.7x10<sup>6</sup><br>4.7% |
| *C. neoformans*<br>**cap10Δ** | 9.6x10<sup>3</sup><br>0% | 2.3x10<sup>5</sup><br>0% | 1.1x10<sup>4</sup><br>0% | 3.8x10<sup>5</sup><br>0% | 1.5x10<sup>4</sup><br>0% | 7.6x10<sup>5</sup><br>0% |
| *C. neoformans*<br>**cap59Δ** | 9.9x10<sup>2</sup><br>0% | 7.9x10<sup>4</sup><br>0.3% | 1.1x10<sup>3</sup><br>0% | 7.6x10<sup>5</sup><br>0% | 2.5x10<sup>5</sup><br>1.8% | 1.0x10<sup>6</sup><br>2.4% |
| *C. neoformans*<br>**JEC20** | 7.9x10<sup>1</sup><br>0% | 3.2x10<sup>4</sup><br>1.8% | 2.4x10<sup>2</sup><br>0% | 2.5x10<sup>3</sup><br>0% | 1.4x10<sup>3</sup><br>0% | 8.6x10<sup>5</sup><br>5.4% |
| *C. neoformans*<br>**JEC21** | 5.8x10<sup>5</sup><br>0% | 8.7x10<sup>5</sup><br>0% | 2.1x10<sup>5</sup><br>0% | 1.2x10<sup>6</sup><br>0% | 7.9x10<sup>5</sup><br>0% | 1.4x10<sup>6</sup><br>0% |
| *C. aspenensis*<br>**DS573** | 8.7x10<sup>5</sup><br>0% | 4.8x10<sup>6</sup><br>0% | 9.8x10<sup>5</sup><br>0% | 2.6x10<sup>6</sup><br>0% | 1.2x10<sup>6</sup><br>0% | 2.3x10<sup>6</sup><br>0% |
| *C. aspenensis*<br>**DS572** | 4.7x10<sup>5</sup><br>0% | 6.1x10<sup>6</sup><br>0% | 2.3x10<sup>5</sup><br>0% | 9.6x10<sup>5</sup><br>0% | 3.2x10<sup>6</sup><br>0% | 4.9x10<sup>6</sup><br>0% |
| *C. aspenensis*<br>**DS570** | 9.1x10<sup>4</sup><br>0% | 6.7x10<sup>5</sup><br>0% | 1.1x10<sup>5</sup><br>0% | 8.9x10<sup>5</sup><br>0% | 9.4x10<sup>5</sup><br>0% | 1.1x10<sup>6</sup><br>0% |
| *C. aspenensis* | 2.1x10<sup>5</sup> | 9.2x10<sup>5</sup> | 1.3x10<sup>5</sup> | 2.4x10<sup>6</sup> | 1.8x10<sup>6</sup> | 1.3x10<sup>6</sup> |
| **DS569** | 0% | 0% | 0% | 0% | 0% | 0% |
| *C. kuetsingii*<br>**MUCL27749** | 2.3x10<sup>4</sup><br>0% | 5.2x10<sup>5</sup><br>0% | 9.8x10<sup>3</sup><br>0% | 9.6x10<sup>5</sup><br>0% | 1.3x10<sup>5</sup><br>0% | 7.7x10<sup>5</sup><br>0% |
| *C. laurentii*<br>**DS620** | 7.9x10<sup>4</sup><br>0% | 9.8x10<sup>5</sup><br>0% | 8.4x10<sup>4</sup><br>0% | 9.7x10<sup>6</sup><br>0% | 1.2x10<sup>6</sup><br>0% | 5.8x10<sup>6</sup><br>0% |
| *C. laurentii*<br>**DS386** | 1.1x10<sup>5</sup><br>0% | 9.6x10<sup>5</sup><br>0% | 9.7x10<sup>4</sup><br>0% | 4.4x10<sup>6</sup><br>0% | 3.8x10<sup>6</sup><br>0% | 1.8x10<sup>6</sup><br>0% |
| *C. laurentii*<br>**DS619** | 1.1x10<sup>5</sup><br>0% | 8.7x10<sup>5</sup><br>0% | 4.4x10<sup>5</sup><br>0% | 8.7x10<sup>6</sup><br>0% | 2.3x10<sup>6</sup><br>0% | 5.4x10<sup>6</sup><br>0% |
| *C. laurentii*<br>**DS621** | 2.4x10<sup>5</sup><br>0% | 9.2x10<sup>6</sup><br>0% | 3.7x10<sup>5</sup><br>0% | 2.2x10<sup>7</sup><br>0% | 3.2x10<sup>7</sup><br>0% | 8.3x10<sup>7</sup><br>0% |
| *C. laurentii*<br>**DS288** | 2.1x10<sup>5</sup><br>0% | 7.4x10<sup>6</sup><br>0% | 2.7x10<sup>5</sup><br>0% | 3.4x10<sup>7</sup><br>0% | 5.6x10<sup>7</sup><br>0% | 9.8x10<sup>7</sup><br>0% |
| *C. uniguttulatus*<br>**MD29** | 1.0x10<sup>4</sup><br>0% | 1.4x10<sup>6</sup><br>0% | 2.3x10<sup>4</sup><br>0% | 7.3x10<sup>5</sup><br>0% | 8.3x10<sup>4</sup><br>0% | 2.4x10<sup>5</sup><br>0% |
| *C. uniguttulatus*<br>**MD26** | 9.1x10<sup>3</sup><br>0% | 8.4x10<sup>5</sup><br>0% | 6.3x10<sup>3</sup><br>0% | 8.7x10<sup>3</sup><br>0% | 8.6x10<sup>3</sup><br>0% | 3.5x10<sup>5</sup><br>0% |
| *C. albidus*<br>**MD22** | 9.0x10<sup>4</sup><br>0% | 9.3x10<sup>5</sup><br>0% | 1.2x10<sup>5</sup><br>0% | 1.4x10<sup>5</sup><br>0% | 7.6x10<sup>5</sup><br>0% | 6.9x10<sup>5</sup><br>0% |
| *C. albidus*<br>**MD41** | 3.2x10<sup>5</sup><br>0% | 8.3x10<sup>6</sup><br>0% | 1.1x10<sup>5</sup><br>0% | 1.4x10<sup>6</sup><br>0% | 1.2x10<sup>6</sup><br>0% | 5.3x10<sup>6</sup><br>0% |
| *C. terrestris*<br>**DS291** | 9.7x10<sup>5</sup><br>0% | 7.6x10<sup>6</sup><br>0% | 4.3x10<sup>6</sup><br>0% | 5.6x10<sup>6</sup><br>0% | 1.2x10<sup>6</sup><br>0% | 8.5x10<sup>6</sup><br>0% |
| *C. terrestris*<br>**DS233** | 1.8x10<sup>5</sup><br>0% | 7.2x10<sup>5</sup><br>0% | 2.9x10<sup>5</sup><br>0% | 2.1x10<sup>6</sup><br>0% | 3.9x10<sup>5</sup><br>0% | 7.5x10<sup>5</sup><br>0% |
| *C. terrestris*<br>**DS234** | 1.7x10<sup>6</sup><br>0% | 8.2x10<sup>6</sup><br>0% | 4.1x10<sup>6</sup><br>0% | 8.0x10<sup>6</sup><br>0% | 2.4x10<sup>6</sup><br>0% | 7.4x10<sup>6</sup><br>0% |
| *C. terrestris*<br>**DS290** | 2.3x10<sup>6</sup><br>0% | 8.2x10<sup>7</sup><br>0% | 4.1x10<sup>6</sup><br>0% | 8.0x10<sup>7</sup><br>0% | 1.4x10<sup>7</sup><br>0% | 7.4x10<sup>7</sup><br>0% |
| *C. terreus*<br>**DUMC177.8** | 7.9x10<sup>2</sup><br>0% | 8.7x10<sup>3</sup><br>0% | 5.4x10<sup>2</sup><br>0% | 8.5x10<sup>3</sup><br>0% | 1.1x10<sup>3</sup><br>0% | 8.7x10<sup>3</sup><br>0% |
| *C. curvatus*<br>**MD110/2009** | 1.7x10<sup>4</sup><br>0% | 2.3x10<sup>5</sup><br>0% | 2.7x10<sup>4</sup><br>0% | 7.9x10<sup>6</sup><br>0% | 1.5x10<sup>6</sup><br>0% | 5.4x10<sup>6</sup><br>0% |
| *C. curvatus*<br>**MD120/2009** | 2.6x10<sup>5</sup><br>0% | 3.1x10<sup>6</sup><br>0% | 4.3x10<sup>5</sup><br>0% | 3.6x10<sup>7</sup><br>0% | 1.2x10<sup>7</sup><br>0% | 4.8x10<sup>7</sup><br>0% |
| *S. cerevisiae*<br>**BY4741** | 2.5x10<sup>4</sup><br>0% | 2.1x10<sup>5</sup><br>0% | 8.6x10<sup>4</sup><br>0% | 1.8x10<sup>6</sup><br>0% | 5.4x10<sup>5</sup><br>0% | 4.3x10<sup>6</sup><br>0% |
| *M. furfur*<br>**CBS 14141** | 3.7x10<sup>3</sup><br>0% | 2.1x10<sup>5</sup><br>0% | 5.3x10<sup>3</sup><br>0% | 3.2x10<sup>4</sup><br>0% | 1.3x10<sup>4</sup><br>0% | 1.4x10<sup>6</sup><br>0% |
| *M. sympodialis*<br>**ATCC 42132** | 2.6x10<sup>3</sup><br>0% | 2.9x10<sup>5</sup><br>0% | 2.7x10<sup>3</sup><br>0% | 1.1x10<sup>4</sup><br>0% | 2.4x10<sup>4</sup><br>0% | 9.1x10<sup>5</sup><br>0% |

\*\* According to standard conditions recommended in protocol for Titan cell induction (except the incubation temperature was set at 30°C), (8)

**Table S4.** Number of cells^A^ and percentage of Titan cells^B^ of two *C. neoformans* var. *grubii* strains depending on different types of FBS and various treatments. The dialyzed FBS (from Gibco) was supplemented with glucose and amino acids (see Materials and Methods for more details). The initial inoculum was 10^3^ cells.

| Tested strain | Fetal bovine serum (name of the company, catalog number) |  |  |  |  |  |  |  |
| --- | --- | --- | --- | --- | --- | --- | --- | --- |
|  | Atlas Biol.<br>F-0500-A |  | Sigma<br>F6178 |  | Gibco (dialyzed) + glucose and amino acids<br>26400-044 |  |  |  |
|  | 48 h incubation at 37°C with 5% CO<sub>2</sub> |  |  |  |  |  |  |  |
|  |  | 15 μM<br>NaN<sub>3</sub> |  | 15 μM<br>NaN<sub>3</sub> |  | 15 μM<br>NaN<sub>3</sub> | inactivated by<br>boiling | inactivated<br>by boiling<br>10 mM AA |
| H99 | <sup>A</sup>6.2x10<sup>5</sup><br><sup>B</sup> 8.3% | <sup>A</sup>3.4x10<sup>6</sup><br><sup>B</sup> 11.6% | <sup>A</sup>5.8x10<sup>5</sup><br><sup>B</sup> 7.9% | <sup>A</sup>4.9x10<sup>6</sup><br><sup>B</sup> 10.9% | <sup>A</sup>6.4x10<sup>7</sup><br><sup>B</sup> 10.8% | <sup>A</sup>4.6x10<sup>8</sup><br><sup>B</sup> 9.9% | <sup>A</sup>2.4x10<sup>8</sup><br><sup>B</sup> 2.3% | <sup>A</sup>9.6x10<sup>8</sup><br><sup>B</sup> 1.2% |
| Btc012Gbc98-1 | <sup>A</sup>2.1x10<sup>6</sup><br><sup>B</sup> 9.6% | <sup>A</sup>6.2x10<sup>7</sup><br><sup>B</sup> 12.2% | <sup>A</sup>4.3x10<sup>6</sup><br><sup>B</sup> 8.2% | <sup>A</sup>3.9x10<sup>7</sup><br><sup>B</sup> 11.7% | <sup>A</sup>9.2x10<sup>7</sup><br><sup>B</sup> 11.7% | <sup>A</sup>5.2x10<sup>8</sup><br><sup>B</sup> 10.2% | <sup>A</sup>9.1x10<sup>8</sup><br><sup>B</sup> 1.3% | <sup>A</sup>9.9x10<sup>8</sup><br><sup>B</sup> 0.8% |
|  | 48 h incubation at 37°C without 5% CO<sub>2</sub> |  |  |  |  |  |  |  |
|  |  | 15 μM<br>NaN<sub>3</sub> |  | 15 μM<br>NaN<sub>3</sub> |  | 15 μM<br>NaN<sub>3</sub> | inactivated by<br>boiling | inactivated by<br>boiling<br>10 mM AA |
| H99 | <sup>A</sup>0.0<br><sup>B</sup> 0% | <sup>A</sup>1.3x10<sup>1</sup><br><sup>B</sup> 0% | <sup>A</sup>0.0<br><sup>B</sup> 0% | <sup>A</sup>2.3x10<sup>1</sup><br><sup>B</sup> 0% | <sup>A</sup>4.5x10<sup>7</sup><br><sup>B</sup> 0% | <sup>A</sup>9.2x10<sup>7</sup><br><sup>B</sup> 0% | <sup>A</sup>7.6x10<sup>8</sup><br><sup>B</sup> 0% | <sup>A</sup>9.4x10<sup>8</sup><br><sup>B</sup> 0% |
| Btc012Gbc98-1 | <sup>A</sup>0.0<br><sup>B</sup> 0% | <sup>A</sup>0.0<br><sup>B</sup> 0% | <sup>A</sup>0.0<br><sup>B</sup> 0% | <sup>A</sup>0.0<br><sup>B</sup> 0% | <sup>A</sup>3.5x10<sup>8</sup><br><sup>B</sup> 0% | <sup>A</sup>8.1x10<sup>8</sup><br><sup>B</sup> 0% | <sup>A</sup>8.5x10<sup>8</sup><br><sup>B</sup> 0% | <sup>A</sup>9.2x10<sup>8</sup><br><sup>B</sup> 0% |

\*Fetal Bovine Serum (FBS)
<sup>A</sup>Number of cells after 48 h incubation (initial inoculum was 10<sup>3</sup> cells)
<sup>B</sup>Percentage of Titan cells after 48 h incubation
AA – ascorbic acid

**Figure S1.**
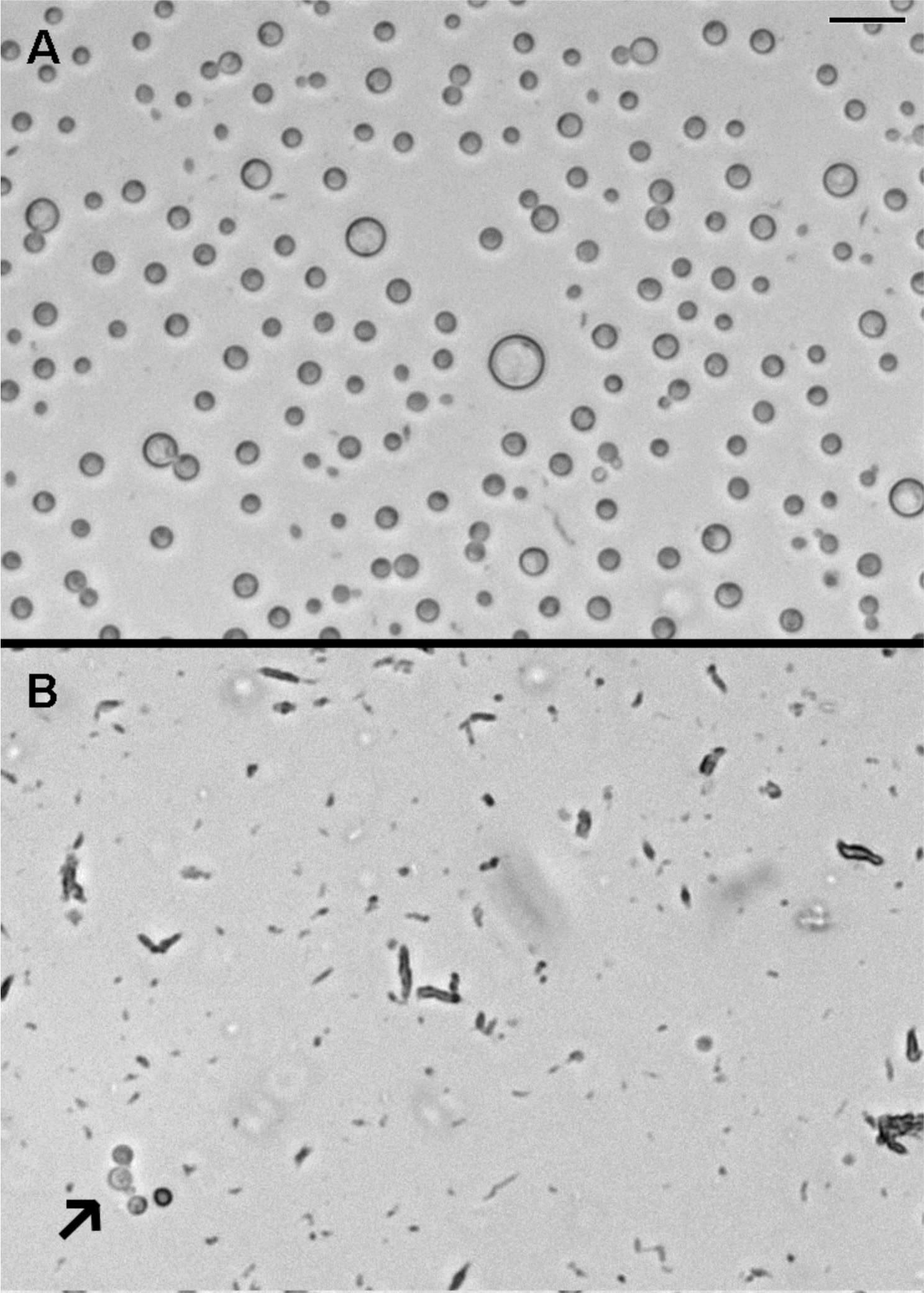
FBS kills *C. neoformans* var. *grubii* and inhibits growth of *C. gattii* upon incubation at 37°C in the absence of 5% CO_2_. *C. neoformans* (strain H99) was incubated under conditions based on Dambuza et al. (8) (A) or identical conditions except 5% CO_2_ was not present during incubation (B). After 48 h, the content of the microplate well was examined under the microscope. Under conditions with 5% CO_2_ (A) cells have proliferated and enlarged Titan cells were formed. Under conditions in the absence of 5% CO_2_ (B) growth inhibition and cell lysis have occurred. The arrow in B indicates a cell rarely detected under these conditions. Bar represents 10 microns.

**Figure S2.**
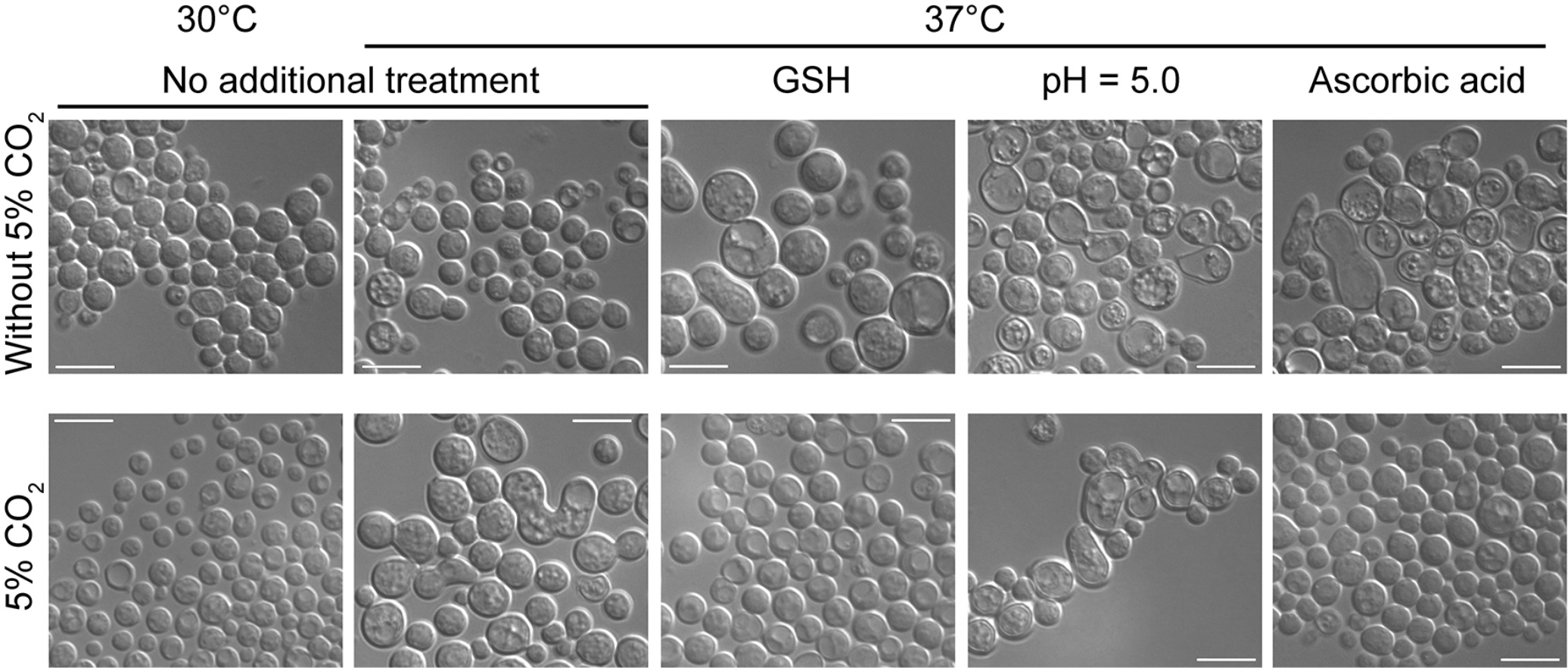
*C. neoformans* var. *neoformans*, strain JEC21 is not capable of forming Titan cells. Cells were subject to titanisation conditions modified from Dambuza et al. (8) as indicated. While some conditions led to cell enlargement, these morphologically changed cells were not round, lacked single large vacuole and a thick cell wall. Bar represent 10 microns.

**Figure S3.**
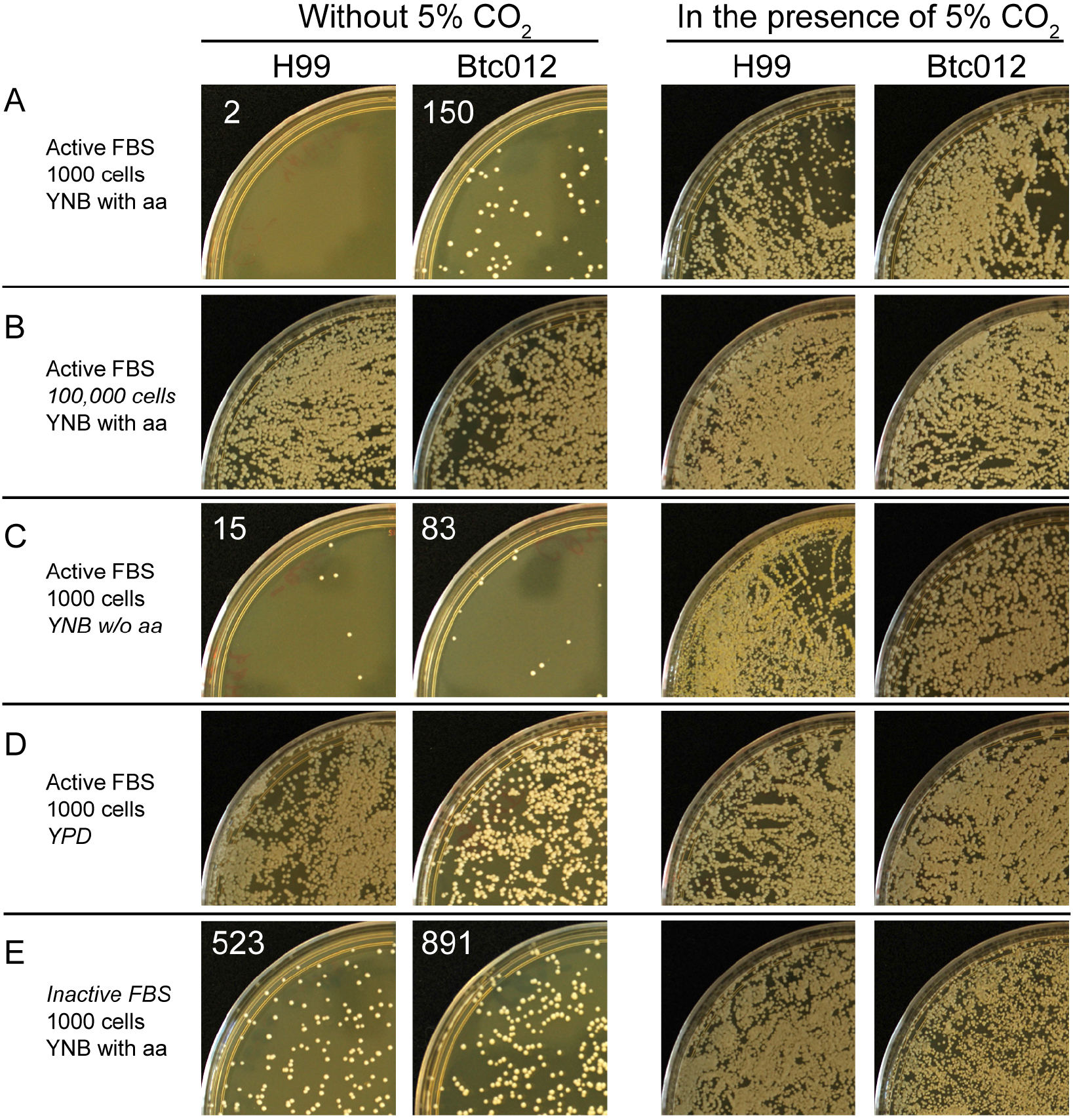
The effects of cell density, YNB type, and heat inactivation of FBS on growth inhibition caused by FBS. The following conditions were tested: (A) Cells were pre-icubated in YNB containing amino acids. ~1000 cells were seeded into the titanisation medium containing 10% of active FBS (Corning 37-075-CV). (B) Same as A, except ~100,000 cells were seeded into titanisation medium. (C) same as A, except YNB without amino acids was utilized for pre-incubation. (D) same as A, except YPD was utilized for pre-incubation. (E) same as A, except heat incactivated FBS (Corning 35-011-CV) was utilized. To test the survival rate after 48 hours of incubation, 100 microliters of the medium (out of 1 ml that was in the well) containing the cells were first diluted 100 times and then spread over the semisolid YPD medium and incubated at 30°C for 2 days. For conditions A, C, and E of the treatment without 5% CO_2_, the entire content of the well was spread on the YPD semisolid media and incubated as described above. The numbers in the top left corners indicate the total number of colonies counted on the plate. Those numbers indicate the number of cells remaining viable out of the original ~1000 cells that were seeded in the wells.

